# Replicative aging is associated with loss of genetic heterogeneity from extrachromosomal circular DNA in *Saccharomyces cerevisiae*

**DOI:** 10.1101/2020.02.11.943357

**Authors:** Iñigo Prada-Luengo, Henrik D. Møller, Rasmus A. Henriksen, Qian Gao, Camilla E. Larsen, Sefa Alizadeh, Lasse Maretty, Jonathan Houseley, Birgitte Regenberg

## Abstract

Circular DNA of chromosomal origin form from all parts of eukaryotic genomes. In yeast, circular rDNA accumulates as cells divide, contributing to replicative aging. However, little is known about how other chromosome-derived circles segregate and contribute to geneticvariation as cells age. We identified circular DNA across the genome of young *S. cerevisiae* populations and their aged descendants. Young cells had highly diverse circular DNA populations, but lost 94% of the different circular DNA after 20 divisions. Circles present in both young and old cells were characterized by replication origins and included circles from unique regions of the genome, rDNA circles and telomeric *Y’* circles. The loss in genetic heterogeneity in aged cells was accompanied by massive accumulation of rDNA circles >95% of all circular DNA. We discovered circles had flexible inherence patterns. Glucose limited conditions selected for cells with glucose-transporter gene circles, [*HXT6/7^circle^*], and up to 50% of cells in a population carried them. [*HXT6/7^circle^*] cells were eventually substituted by cells carrying stable chromosomal *HXT6 HXT6/7 HXT7* amplifications, suggesting circular DNA were intermediates in chromosomal amplifications. In conclusion, DNA circles can offer a flexible adaptive solution but cells lose genetic heterogeneity from circular DNA as they undergo replicative aging.

## INTRODUCTION

In the unicellular eukaryote model, *Saccharomyces cerevisiae*, aging is determined by the number of replicative cycles a mother cell undergo before dying (1). The asymmetric division of yeast causes an accumulation of factors in the mother cell such as protein aggregates (2, 3) and circular DNA from the rDNA locus, also known as ERCs or [*rDNA^circle^*] (4). Accumulation of circular rDNA is associated to a massive increase in ribosomal rRNA levels (5) and the accumulation can reduce the number of divisions a cell can undergo by 40% (4). Replicative aging is not specifically associated with circular DNA from the rDNA locus but rather it appears to be the accumulation of circular DNA in general that causes a shorter lifespan of the mother cell (4).

The segregation and accumulation of circular rDNA has been extensively studied (4–10). However, our knowledge about segregation of other circular DNA species of chromosomal origin is based on a few model circles with a small range of genetic markers (4, 6, 7, 11). Therefore, it is not clear whether segregation kinetics, inferred from a small number of model circles, can be generalized to circular DNA formed from other parts of the genome. This is important to establish since circular DNA, derived from other parts of the genome, might naturally contribute to aging and because amplifications of genes on circular DNA can provide strong phenotypic effects to the cell carrying them (12–16).

Circular rDNA from the rDNA array on chromosome XII carries a consensus sequence for replication (ARS) for each rDNA-repeat unit (17). This can likely ensure replication of a formed [*rDNA^circle^*] in mitosis but not faithful segregation between mother and daughter cells (6, 7, 11). The asymmetric segregation of replicated rDNA is suggested to be the cause of accumulation of [*rDNA^circles^*] in mother cells (4). It was shown that circular rDNA mis-segregate with an estimated retention frequency above 0.99 per circle towards the mother cell (10). The segregation bias can explain accumulation of replicating circular rDNA though the amount of circular rDNA might also be affected by increased formation rates in aging cells (18), available nutrients and polymerase I (POLI) activity (9), as well as the length of the rDNA array (9).

The fate of circular DNA from other parts of the genome has primarily been studied through plasmids with the marker genes *LYS1* and *URA3* (4, 6–8, 11). Circular DNA without centromeres have segregation biases that are similar to those for rDNA (6, 11) and two models have been suggested to explain their tendency to segregate towards the mother cell. In the active retention model, rDNA and non-rDNA circles are actively retained in the mother cells by the nuclear pore complexes via the TREX-2 and SAGA complex, which are suggested to tether the circular DNA to nuclear periphery of the mother cell (8). The alternative model, known as the diffusion model, suggests that segregation of non-rDNA circles is determined by passive diffusion from the mother nucleus to the daughter nucleus (6). The nuclear geometry restricts diffusion of circular DNA from mother to daughter under mitosis and thereby lead to a segregation bias (6). The passive model is based on *LYS2* circles with and without replication origin that distribute randomly across the nucleus rather than being associated to the nuclear envelope (6). The different observations of circle distributions in the nucleus during mitosis, suggest that more than one mechanism for mis - segregation might exist.

Other genetic elements on circular DNA can ensure more equal segregation between mother and daughter cells. Circles with replication origins and centromeres are mitotically stable through their anchoring to the mitotic spindle and deviation from a 1:1 segregation is only expected when centromeric circles fail to replicate (11). Anchoring of circular DNA to telomeres through telomere binding proteins also ensures near equal segregation between mother and daughter cells (6, 19).

Recent studies have shown that circular DNA form from all parts of eukaryotic genomes (20–22). While the majority of the circular DNA are smaller than 1 kilobase (kb) (20, 22, 23), some circular DNA are large enough to fully span transposons (24), centromeres, genes, and replication origins (13, 20, 22, 24–26). Hence, circular DNA can acquire all the genetic elements required for the replication and propagation of the genetic material present on them. Yet, a global overview of how circular DNA segregates and affects aging is still missing.

To obtain a global overview of how circular DNA segregates, we have applied the yeast Mother Enrichment Program (MEP) to obtain young and the corresponding aged cells (27) and combined it with our methods for purification and mapping of circular DNA (28, 29). This has allowed us to dissect the inheritance pattern of circular DNA while *S. cerevisiae* cells undergo replicative aging. To get a robust measure of the different circular DNA species present in young and aging cells, we have developed novel methods for mapping of circular DNA from both uniquely mappable and repetitive regions of the *S. cerevisiae* genome. Besides identifying circular DNA with different segregation patterns, we have adapted methods for quantifying the copy number levels of the detected circular DNA. This has allowed us to cluster circular DNA species and search for consensus elements that determine their inherence patterns. We find that most circular DNAs are lost from aging cells, while circular DNA with replication origins are prone to be maintained in cells as they age but only a few of these maintained circular DNA accumulate in aging cells.

## MATERIAL AND METHODS

### Cells

Strains for preparing circular DNA datasets were: UCC5185 (diploid) with the genotype S288C *MAT*a/*MAT*α *ade2::hisG/ade2::hisG his3/his3 leu2/leu2 LYS2/lys2 ura3Δ0/ura3Δ0 trp1Δ63/trp1Δ63 MET15/met15Δ::ADE2 hoΔ::PSCW11-cre-EBD78-NATMX/hoΔ::PSCW11-creEBD78-NATMX loxP-UBC9-loxP-LEU2/loxP-UBC9-loxP-LEU2 loxP-CDC20-Intron-loxPHPHMX/loxP-CDC20-Intron-loxP-HPHMX* (27). Cells carrying [*GAP1^circle^*] in the CEN.PK background (G1 and G2) served as the positive control for circular DNA (13).

### Separation of progeny, aged and young populations

MEP 2-5 samples were grown overnight in YPD media (2% peptone, 1% yeast extract, 2% glucose) at 30°C with shaking at 200 rpm to mid-log prior to labelling. 6-10 MEP samples for datasets 6-10 were transformed with an empty plasmid for quantification purposes (pRS316) then individual colonies picked and grown overnight in synthetic complete medium minus uracil to mid-log. Media components were purchased from Formedium and media was sterilized by filtration. [*GAP1^circle^*] strains (13) were propagated in minimal medium with L-glutamin as sole nitrogen source to sustain the [*GAP1^circle^*] (2% Dextrose, 0.16 % yeast nitrogen base without ammonium sulfate and amino acids, 0.34 mM L-glutamine). Upon initiation of the experiment, cells were transferred to nonselective conditions, YPD, for 5 generations.

For biotin labelling, 1×10^7^ cells were harvested by centrifugation (15 s at 13,000 g), washed twice with 500 µL PBS and re-suspended in 500 µL of PBS containing ∼3 mg/mL Biotin-NHS (Sigma B1022). Cells were incubated for 30 min on a rotating wheel at room temperature, washed once with 125 µL PBS and re-suspended in YPD. One sample of 0.25×10^7^ cells was frozen on liquid nitrogen (N_2_) at this point (B0). For datasets 2-5, 0.25×10^7^ cells were inoculated in 12 mL YPD in a 25 mL flask and grown for 8 hours at 30°C with shaking before harvest after 5 doublings by centrifugation and freezing on N_2_ (progeny population). 0.5×10^7^ cells were inoculated in 500 mL YPD in a 1 L flask containing 1 mM β-estradiol in a 500 mL flask and grown for 48 hours as above before harvest (BA). For datasets 6-10, 0.5×10^7^ cells were inoculated in 50 mL YPD in a 100 mL flask and grown for 8 hours at 30°C with shaking, followed by harvesting by centrifugation and freezing on N_2_ (progeny population). 0.25×10^7^ cells were inoculated in 150 mL YPD containing 1 mM β-estradiol in a 500 mL flask and grown for 24 hours as above before harvest (aged population).

For cell purification, gradients (1 for progeny population or 2 for aged population) were formed by vortexing 1.16 mL Percoll (Sigma P1644) with 42 µL 5 M NaCl and 98 µL water in 2 mL tubes and centrifuging 15 min at 15,000 g, 4°C. Cells were defrosted and washed 1x with 1 volume of cold PBS + 2 mM EDTA (PBSE) before resuspension in ∼250 µL cold PBSE per gradient and layering on top of the pre-formed gradients. Gradients were centrifuged for 20 min at 1,000 g, then the upper phase and brown layer of cell debris removed and discarded. 1 mL PBSE was added, mixed by inversion and centrifuged 1 min at 2,000 g to pellet the cells, which were then re-suspended in 1 mL PBSE per time point (re-uniting samples split across two gradients). 50 µL streptavidin microbeads were added (Miltenyi Biotech 1010007) and cells incubated for 30 min on a rotating wheel at room temperature. Meanwhile, 1 LS column per sample (Miltenyi Biotech 1050236) was equilibrated with cold PBSE in 4°C room. Cells were loaded on columns and allowed to flow through under gravity, washed with 8 mL cold PBSE and eluted with 1 mL PBSE using a plunger. Cells were re-loaded on the same columns after re-equilibration with ∼500 µL PBSE, washed and re-eluted, and this process repeated for a total of three successive purifications. 50 µL cells were set aside for quality control, while the remainder were pelleted by centrifugation and frozen on N_2_ (progeny+ and aged population fractions). For the progeny fraction, the initial flow-through of unbound cells from the first column was also collected and passed through a second column to deplete any remaining biotin-labelled cells to form the progeny-fraction.

### Cell counting

Three different methods have been used to determine the precise number of cells in different experiments. The *Counting chambers* method. In this method 2 µL of a cell sample in an appropriate dilution was loaded onto the counting chamber (Brand, Cat. No.: 719005). The number of cells was then counted in one chamber, after which the total number of cells per µL was calculated by this formula:

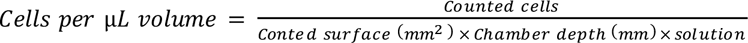

At least five different chambers were counted in order to minimize variation in the counted number of cells. *Hemocytometer:* After cell batch production, each cell sample were pelleted, fixed in 70% ethanol and counted in a hemocytometer after rehydration in PBS. Approximately 1E6 cells were pelleted and frozen on liquid nitrogen (N_2_).

### Circular DNA purification and sequencing

Circular DNA was purified from young, aged and progeny subpopulations of 10^6^ cells with the Circle-Seq protocol (20) with minor changes: Cell membranes were disrupted by 10 units zymolyase (USBiological) for 1.5 hour at 35°C. Zymolyase was inactivated at 60°C for 5 min. To assist sufficient circular DNA precipitation after column chromatography, samples were incubated at -20°C for 45 min and then centrifuged at 9788 × g for 30 min at 2°C, followed by 70 % ice-cold ethanol wash and re-centrifugation at 9788 × g for 5 min at 2°C. The air-dried DNA was dissolved in 19 µL of sterile water and treated with Swa1 in two cycles (2×2 FastDigest units, Thermo Scientific), each cycle of 1 hour at 37°C (total volume 25 µl) and heat-inactivation for 15 min at 65°C. To remove all linear chromosomal DNA, four units of exonuclease (Plasmid-Safe ATP-dependent DNase, Epicentre) was added to the reaction at 37°C (reaction volume 40 µL) and fresh 2.5 units DNase was added every 24-hours with additional ATP and buffer. After digestion for six days, enzymes were heat inactivated for 30 min at 70°C. The liquid (∼ 47 µL) was evaporated to half volume on a vacuum centrifuge (MAXI-DRY LYO, Heto) and 10 µL of the volume was treated with REPLI-g Mini (Qiagen) for 40 hours.

Two different set of plasmid spike-mixtures were used as internal controls. We used a mixture containing the following plasmids to cell ratio pUG72 (1:1000), pUC19-yEGFP (1:10,000), pSH63 (1:1,000), p4339 (1: 10,000), YGPM25009_18kb (1:100), YGPM3k20_chrV_26kb (1:100), pBR322 (1:100) and pRS316 (endogenously maintained) for the 2-5 populations. For the 6-10 populations, we used the following plasmids to cell ratio pUG72 (1:50), pUC19-yEGFP, (1:500), pSH63, (1:500), p4339 and PRS316 (1:500) and pRS316 (endogenously maintained). pUG72 and pUC19-yEGFP were added when cells were lysed and plasmids pSH63 and p4339 were added after exonuclease treatment. Phi29 amplified circular DNA was subsequently fragmented by sonication and the output was sequenced by Illumina (HiSeq 2500) paired-end sequencing with read lengths of 2×141 bp for the 2-5 samples and 2×75 bp for the 6-10 samples. Linear DNA content was analyzed in samples during the exonuclease time course using the following standard PCR methods and primers: *ACT1* 5′-TGGATTCTGGTATGTTCTAGC-3′ and 5′-GAACGACGTGAGTAACACC-3′ and quantitative PCR (qPCR) primers 5′-TCCGTCTGGA TTGGTGGTTCTA-3′ and 5′-TGGACCACTTTCGTCGTATTC-3′. Exonuclease treated total DNA was only amplified by Phi29 when all linear *ACT1* was found to be depleted.

### Quantitative PCR

Quantitative PCR (qPCR) was performed on a QuantStudio 7 PCR system (Applied Biosystem) with Power SYBR® Green PCR Master Mix (Life Technologies, Carlsbad, CA, USA) using the following program: 1 cycle 95°C, 10 minutes; 40 cycles of 95°C, 15 seconds and 60°C for 60 seconds and finally 1 cycle 95°C, 15 seconds and 60°C 15 seconds to assess reaction specificity (melting curve). Primers for quantification of [*rDNA^circles^*] were: 5’-CTTCTTCCCAGTAGCCTCATCC-3’ and 5’-TGAACAGTGAACAGTGGGGAC-3’; [*CUP1-1^circles^*]: 5’-CTGGCAAGTAGAAAGGAACACC 3’ and 5’- TCTCCGATACCTGCCTCTATTG-3’: [*GAP1^circles^*]: 5’-TGTTCTGTCTTCGTCACCGC-3’; 5’- GGCTCTACGGATTCACTGGC-3’; *ACT1*: 5’-TCCGTCTGGATTGGTGGTTCT-3’ and 5’-TGGACCACTTTCGTCGTATTC-3’. All qPCR reactions were performed in quadruplicates measurements and experiments were done minimum twice.

### Genotyping of, [Y’telomeric^circle^], [ENA5 ENA2 ENA1^circle^], [CEN1^circle^] and [GAP1^circle^]

PCR was carried out according to standard procedures. The following inverse PCR primers were used for detection of circular DNA from: [*ENA5 ENA2 ENA1^circle^*], 936 5’-ACTTGTGCCTATAAATTCAGCGG-3’ and 937 5’-GTACTTCTACTACAATCCATACAGAAG-3’; [*CEN1-1^circle^*], 917 5’ ACGAAGACTGAACTGGACGG 3’ and 918 5’ ACGAAGACTGAACTGGACGG 3’ and [*GAP1^circle^*], 961 5′AACTTTGGGGATTCGGAGTTC3′ and 962 5′-TTGATTACTTGACCCAGAGCG-3′. The [*Y’telomeric^circle^*] was amplified with primers that were general for telomeric Y’-elements 892 5’-GATACGATACTTTCTGGC-3’ and 893 5’- GATGGTGTTAGACAAGGC-3’ providing a 5.4 kb product as well with primer 892 and a TEL4R TEL12R TEL15R specific primer 895 5’-CCCAACATAGAAAACAGAAGG-3’. Purified PCR products were next Sanger sequenced for assembly of the [*Y’telomeric^circle^*] sequence. Sequencing was done with 893, 895, 808 5’-ACCGACATTAACAAAGAGTCG-3’; 842 5’-CTGGATAGCGATGATGTTCC-3’, 896 5’-CCTTCTGTTTTCTATGTTGGG-3’, 897 5’-AGTTGAGTTTCCGTGTCC-3’, 898 5’-TGGTACCTTCTTCTGATGG-3’, 899 5’-GTAACAAATCGAGACATTTCG-3’, 900 5’- ATGGCGTTACTGATTTATACG-3’, 901 5’-TGACACAGTGGAACTGATAG-3’. *Y’* from TEL4R, TEL12R and TEL15R are completely identical, and could therefore not be separate by sequencing. These circles contained the entire telomerase encoding Y’-element. PCR reactions were carried out with 0.2 ng of template from Phi29 amplified sample or 4 µL of template from 130-h exonuclease-treated sample using standard conditions.

### Read alignment

We demultiplexed the reads allowing zero mismatches in the barcodes and we checked the quality with FastQC(30). We used the *S. cerevisiae* reference genome (S288C-R64-2-1) and the plasmids included in the Circle-Seq procedure as the alignment reference. Reads were aligned using BWA-MEM (v0.7.17-r1188) (31) with the –q option, leaving the rest of the parameters on their default settings. Samples with a mapping percentage lower than 80% were realigned reducing the seed (-k: 13). We used SAMtools (v1.9) (32) in all our downstream BAM processing steps (Supplementary Figure S1).

### Circular DNA identification

We used Circle-Map (v.0.9) package(29) to identify circular DNA. To detect circular DNA from the alignable regions, we first extracted all the reads with variation using the sub-package Circle-Map ReadExtractor. Then, we executed the Realignment submodule with all the parameters set to default. We obtained a robust circular DNA identifications by iteratively merging the circles that reciprocally overlapped by 50%. Then, we removed the circular DNA that did not contained >4 supporting reads and a sequencing coverage of 90%. We evaluated the effect of the sequencing depth on number of circular DNA by subsampling (setting SAMtools seed to 2) reads on the BAM files at intervals of 5% of reads, and executing the Circle-Map Realign pipeline as described above.

To detect Circular DNA from repetitive regions of the genome we identified reads whose 2sd highest alignment score was >= 80% of the 2st highest alignment score, with edits < 3. For every highest score – 2sd highest score pair, Circle-Map reported the circular DNA using the leftmost and rightmost alignment position of the pair. Then, circular DNA with a read support lower than 20 were removed. To avoid redundant circular DNA, circular DNA reciprocally overlapping by 80% were merged. Then, we removed intervals with a coverage less than 80%.

### Saturation curves

We evaluated the effect of the sequencing depth on number of circular DNA by subsampling (setting the SAMtools random seed to 2) reads on the raw BAM files at intervals of 5% of reads, and executing the Circle-Map Realign pipeline as described above.

### Genomic feature annotation

We annotated genomic features on circular DNA using bedtools (v2.27.1). We downloaded the features from the Saccharomyces Genome Database (Genome Version R64-2-1). Altogether, the following features were annotated: replication origin, gene names (symbols) and ORFs (location holders), centromeres, introns, solo LTR elements, LTR transposons (Ty), telomeric elements, rRNA genes and tRNA genes. We annotated experimentally validated origins of replication from OriDB database (33) transforming the coordinates to the SacCer3 version using the UCSC liftover tool (34).

### Classification of inheritance patterns

Circular DNA found in both young, progeny and aged population were determined by intersecting the chromosomal coordinates. We allowed a maximum distance of ± 300bp in the coordinates of circular DNA from alignable regions and 400bp for those from repetitive regions. We classified the circular DNA to class I if we found it all the subpopulations a sample; class II (in young and aged), class III (in young and progeny) and class VI (in aged and progeny); class IV (only in the young), class VII (only in the progeny) and class V (only in the aged). We applied two post -processing steps. If a circular DNA was classified to different, we assigned the circular DNA to the highest order of representation (I > II, III & IV > V, VI & VII). Then, circular DNA found in two classes of the same order of representation were considered ambiguous and we removed them from the further analysis. To discard whether the low number of intersections between the young and aged samples was produced by our filters to the data, we intersected all the circular DNA classified as lost with all the detected circles in the aged samples using the same distance criteria as mentioned above but without applying any read or coverage filters to the data.

### *De novo* mutation rate

We calculated the *de novo* mutation rates by dividing the number of circular DNA detected in the 2-5 aged yeast population by the cells recovered in the 4 aged populations multiplied by the number of divisions the aged cells went (20 divisions).

### Circular DNA quantification

We counted the unambiguously aligned reads (quality >= 20) to every circular DNA across samples. We used R (v3.5.1) (35) and DESeq2 (v1.22.2) to normalize the read counts with the median of ratios method followed by log10 scaling. We analyzed circular DNA detected in at least two samples using principal component analysis, (PCA, with scale and center parameters set to true) and hierarchical clustering using the pheatmap (v1.0.12) package. We used the Manhattan distance for row and column clustering and the average linkage algorithm for defining the clusters. To calculate log fold-change distributions, we combined the circular DNA count matrices from samples 2-5 and 6-10 and we computed log10 fold changes between the aged and the young populations for all the circles that were detected on the young datasets. We used Gviz (v1.24.0) (36) to create the coverage plots of the selected circles.

Alternatively, we used the spike in plasmids in samples 6 to 10 to quantify the circular DNA levels. We first counted the number of reads aligned to the plasmids and the detected circular DNA (quality >= 20) and to the telomeric Y’ and rDNA region (no mapping quality filters due to its tandem duplication structure). Then, we divided the number of reads aligned to the rDNA region by the sum over all reads aligned to plasmid (pUC19_yEGFP, pSH63, p4339 and pRS316).

### Genomic feature overrepresentation

We tested the overrepresentation of the annotated genomic features (OriDB, centromere, long_terminal_repeat, telomere, LTR_retrotransposon and intron) and 80 cis acting elements binding sites (37) between the class I & II and class III & IV circles. To define if a circular DNA carried a cis acting element, we scanned the sequence of the circular DNA for an exact match of the motif, with the exception of the origins of replication were we allowed for non-exact overlaps due to the lack of resolution. Then, for every feature, we used a two tailed Fisher test (38). We independently corrected the p-values for the cis acting elements and the genomic features using the Benjamini-Hochberg method.

### Genotyping of cells in prolonged glutamine-limited chemostat cultures

Cells were isolated from a CEN.PK glucose limited chemostat (CEN.PK wt *MAT*α G6, as in (39)) at generations 0, 35, 71, 82, 106, 124 and 135. DNA was purified from 96 clones from each time point, by growing cells in microtiter plates in 300 µl YPGal (1% Yeast Extract; 2% Peptone, 1% Galactose). When outgrown, cells were washed with 0.9 M sorbitol and 0.1 M EDTA and exposed to Zymolyase (Zymo Research) at 37 C° until a majority of cells burst when exposed to hypo-osmotic stress in water. Cells were then pelleted and suspended in 125 µl P1 buffer (Qiagen), that was then mixed with 125 µl P2 buffer (Qiagen) and 175 µl of N3 buffer (Qiagen). Denatured cell debris was pelleted by centrifugation and 200 µl supernatant transferred to 200 µl isopropanol for precipitation of DNA with 30 minutes, 4000 rpm centrifugation. Pelleted DNA was finally washed in 70% EtOH and suspended in H_2_O. The following PCR primers were used for genotyping: 654 5’-TACCGAGGTGAGCCCTGC-3’; 655 5’-TCCGTCAGAGGCTGCTACG-3’; 656 5’ CTGACTTCTTCCCACTTTGC-3’; 657 5’-GACAATGGAGAGCAAATGGG-3’; 749 5’-TCTTCAGTTTGACCAGCAACC-3’ and 751 5’-TCGCTTCTCAACAAGATTTGC-3’. A 7.5 kb PCR product on genomic DNA with 655 and 656 and no PCR product with 654 and 655 was indicative of wt *HXT6 HXT7*; a 2.1 kb PCR product on genomic DNA with 656 and 657 was indicative of a *HXT6/7* deletion; and a 2.7 kb PCR product with primers 654 and 655 was indicative of an amplifications of the *HXT6 HXT7* locus. Finally, [*HXT6/7^circle^*] was detected in samples enriched for circular DNA by BamHI digest and plasmid-safe ATP DNAse (Epicenter E3101K) treatment of linear DNA (BamHI does not cut the *HXT6 HXT7* locus). Samples were found to contain [*HXT6/7^circle^*] if a 2.1 Kb product was obtained from the exonuclease treated DNA with the 656 and 657 primer set and no products with the chromosomal control with primers 749 and 751. Selected PCR products were sequenced by Sanger sequencing to confirm genotypes. Genotyping was repeated on the independent glucose limited chemostat culture (CEN.PK wt MATα G7 described in(39)).

### Statistical analysis

We used the Wilcoxon test (R v3.5.1) to test if the medians were different between groups.

## RESULTS

### Genome-wide circular DNA detection in young and aged cells

To determine the fate of circular DNA as *S. cerevisiae* age, we compared circular DNA across the genome of yeast populations when young and aged. This was done in 4 replicate populations, each of which was sampled when “young” and also used to produce two separate subpopulations: 1) an “aged” subpopulation of purified mothers that had aged for 48 hours (∼15-25 cell divisions) through the MEP (27) that allows only mother cells to divide (Figure 1A) and 2) a “progeny” subpopulation that was allowed to produce progeny for 5 cell divisions and harvested without further purification (Supplementary Table S1).

**Figure 1.**
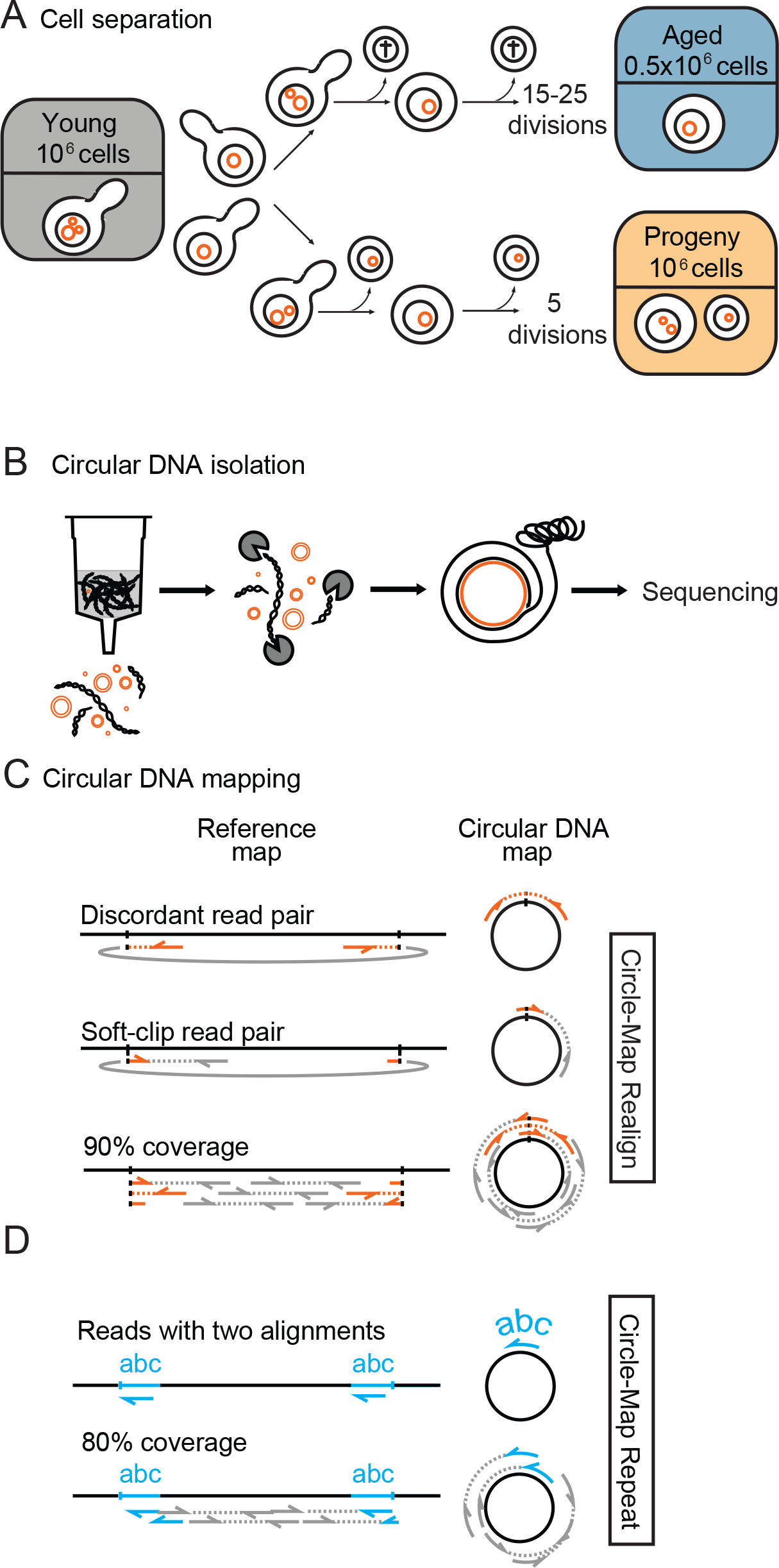
Schematic overview of aged and progeny cell separation using the Mother Enrichment Program, circular DNA purification and mapping. (*A*) Aged, progeny and young cell separation from four populations of yeast cells (named 2, 3, 4 and 5). The number of cell divisions in aged cells was based on bud-scar counts and the number of cell divisions in the progeny population was based on population size that grew from 10^6^ to 32×10^6^ when harvested. Only 10^6^ cells were harvested from each of the progeny populations and thereby only 3.3% of the cells in a population. The aged populations contained on average 7.3×10^5^ cells (S.D: 1.9×10^5^). (*B*) Circular DNA purification and sequencing using the Circle-Seq method. (*C*) Circular DNA detection from the mappable parts of the genome using structural variant reads (soft-clipped, discordant and split reads in orange, concordants in grey) and circular DNA read coverage >90%. (*D*) Circular DNA detection from repetitive regions of the genome using reads with a high scoring suboptimal alignment position (reads with suboptimal alignments in blue, reads without suboptimal alignments in grey) and circular DNA read coverage >80%.

We next purified and sequenced circular DNA from each subpopulation by removal of linear DNA (Figure 1B and Supplementary Figure S2), rolling circle amplification and short-read paired end sequencing as previously described (20). We developed two algorithms to map the origin of circular DNA in the *S. cerevisiae* genome, Circle-Map Realign and Circle-Map Repeats (Figure 1C-D). The first detects circles from the unique regions of the genome by probabilistic alignment of reads that span the junctions of circles (soft-clipped and discordant reads) (29). Using more than four of such junction reads and a read coverage > 90% as cut-off for confident circle detections, we detected 404 unique circular DNA over the 4 populations (subpopulation median: 26.5, range: 9-105) (Supplementary Table S2). The cut-off was found to be the optimal number of breakpoint reads and coverage for a robust recording of circular DNA (Supplementary Figure S3). The second algorithm, Circle-Map Repeats, detects circular DNA from repetitive regions of the genome (Figure 1D) such as those from the rDNA array, LTR retrotransposons and the *HXT6 HXT7* locus (4, 13, 20). We detected 51 repetitive circular DNA (Supplementary Table S2) of which many came from loci known to undergo circularization such as rDNA, *CUP1 CUP2* and *ENA1 ENA2 ENA5* (4, 20). All in all we found 51 + 404 circular DNA species in the 4 populations of young, progeny and aged cells. The experiments were repeated with another four populations where we found 6 repetitive and 61 unique circular DNA (Supplementary Figure S4, Supplementary Table S2).

### Young cells have larger genetic heterogeneity from circular DNA than aged cells

We next classified the circles found in the young cells into five classes (Figure 2A): I) circles found in all populations, II) circles found in both young and aged populations, III) circles found in the young and progeny but not in the aged, IV) circles lost from the population and V) circles formed *de novo*. Our classification revealed that 93.7% of the circles in the young cells (Figure 2B, class IV) were not detected when cells aged, suggesting that the majority of the circles present in young cells were lost as cells aged. To determine if the low number of circles from young subpopulations recurring in aged subpopulations was affected by sequencing, we tested the impact of sequencing depth by subsampling a fraction of the total alignments, mapping and counting of the number of circle species in the subsample (Figure 2C). We found that all circular DNA species were captured at 20-30% of the total reads present in our sequencing datasets (Figure 2C and Supplementary Figure S5). Furthermore, to exclude that circles in the aged populations were filtered out due to our cut -off thresholds, we reanalysed the overlap between the young and the aged populations without applying split-read and read coverage filters (Figure 1C-D) to the aged samples. This analysis gave no read evidence for circular DNA in aged cells for 96.5% of the class IV circular DNAs found in the young cells (e.g. Figure 2D). Hence, our data shows that we detected the majority of circular DNA present in the yeast populations, and that a greater sequencing depth would most likely not have led to detection of more circles in any of the young, progeny or aged subpopulations. Besides class IV circ les that were only found in young cells (Figure 2D), we also found circular DNA that was only in the progeny and/or in the aged cells. These circles are likely to be products of *de novo* circularization events (Figure 2A, class V, aged mothers) that had occurred during the 15-25 divisions the cells underwent after being separated from the young subpopulation, allowing us to estimate the median DNA circle mutation-rate to 2.03×10^−6^ mutations/cell division (range: 7.75×10^−7^ – 3.9×10^−6^). Taken together, these results suggest that the majority of DNA circles present in young cell populations were lost as cells underwent replicative aging.

**Figure 2.**
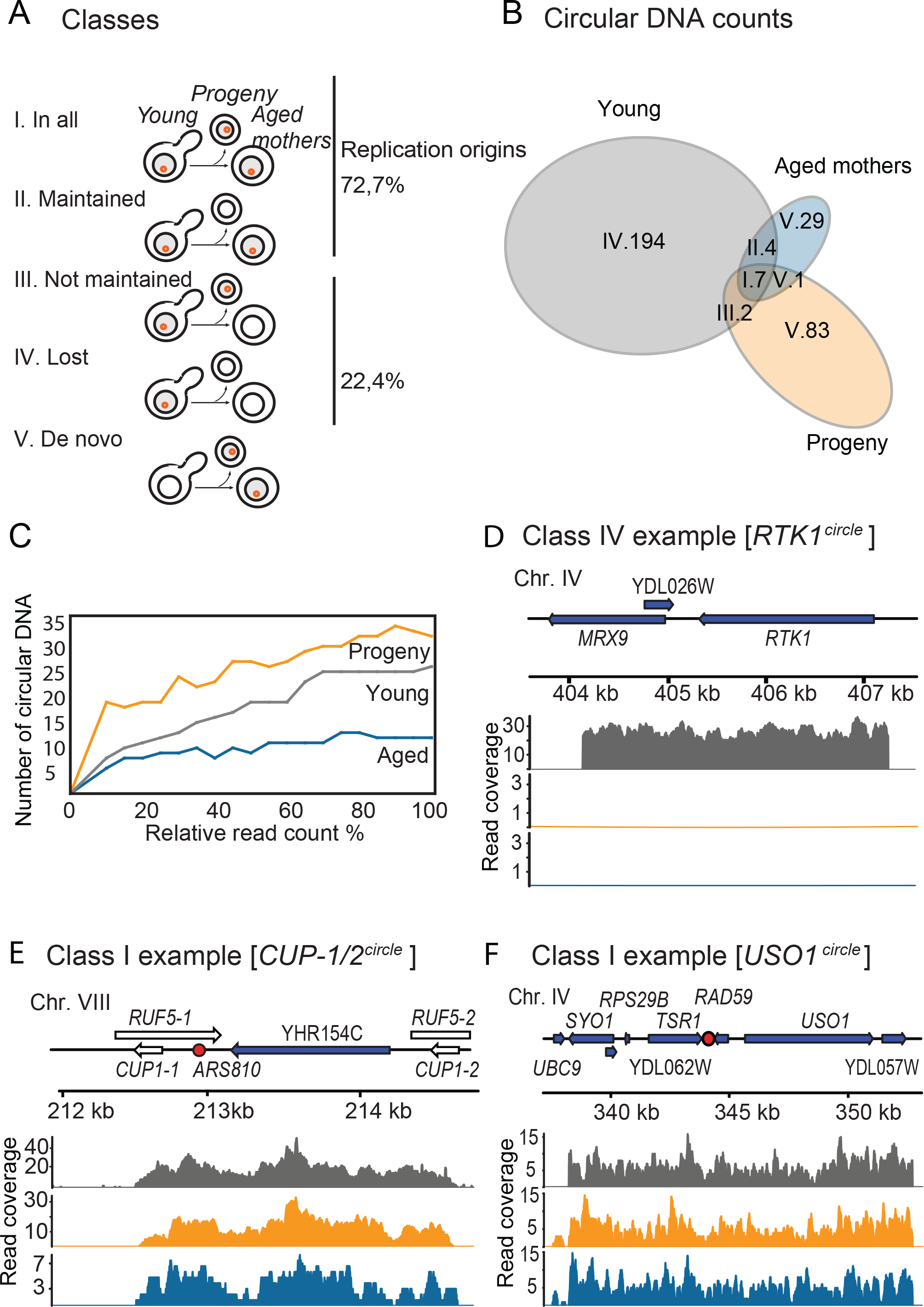
Segregation patterns of circular DNA in yeast cells. (*A*) Schematic overview of the different segregation patterns followed by circles found in the 2-5 young populations as cells divide. (*B*) Group-size aware Venn diagram displaying the segregation of the different DNA circles found in the young, progeny and aged yeast populations. (*C*) Saturation plot representing the circular DNA counts against the percentage of reads sampled from the entire alignment file for the sample 2. (*D-F*) Circular DNA read coverage plots for young (grey), progeny (orange) and aged (blue) samples. The genomic features present in the circular DNA are shown above the coverage plots (repetitive elements shown in white, genes shown in blue and replication origins shown in red). (*D*) Read coverage plot of the [*RTK1^circle^*], a class IV circle. (*F*-*E)* Read coverage of the [*CUP1/2 ^circle^*] and [*USO1^circle^*], both classified as class I.

### Circles present in both young and aged cells are characterized by replication origins

To find traits that could explain why some circular DNA persist in a population and others not, we screened for genetic features on circles maintained in the yeast populations as cells underwent replicative aging. We found that the recurring class I and II circles originated from repeat rich regions of the genome such as the *HXT6 HXT7,* rDNA, and *CUP1-2* loci (Figure 2E, Supplementary Table S2). Still, six out of eleven circles found on this class were from non-repetitive genomic regions (Figure 2F, Supplementary Table S2), and our data indicates that there is no significant enrichment for circles from repetitive regions in the group formed by class I and class II circles (Two sided Fisher’s exact test, n=207, *adjusted p-value =* 0.09). One significantly enriched genomic feature in the class I and II circles were replication origins, found in 8 of the 11 recurring circular DNA while only 44 out of 196 of the class III+IV circles carried origins of replication (Figure 2A and Supplementary Figure S6; Two sided Fisher’s exact test, n=207, *adjusted p-value =* 0.006). We also tested if size or other genomic sequences could explain maintenance of circles in cells as they aged. However, neither size of circular DNA was found significantly overrepresented in the group formed by class I and II circles (Two sided Mann-Whitney U test, n=207, *p-value =* 0.08) nor were the 80 different transcription factor binding sites (Supplementary Table S3)) as well as sets of genomic features such as LTRs, telomeres and centromeres (Supplementary Table S4). Taken together, these results suggest that replication origins are an important feature for maintenance of circular DNA as mother cells divide, as recently shown using *CUP1* as a model (40).

### Recurrence of circular DNA from direct repeat loci and loss of a CEN circle

Among the circular DNA identified, we found circular DNAs that did not follow the inherence patterns expected from their genetic features. These were circles from direct repeats, telomeric Y’-circles and centromeric circles (Figure 3). An example of a circular DNA from a direct repeat locus was the [*ENA^circle^*], which did not carry a replication origin but still occurred recurrently in different subpopulations (Figure 3A, circular DNA diagram). Consequently, we expected [*ENA^circles^*] would fail to be maintained in the population as they lack replicative capacity. [ *ENA^circle^*] forms by recombination (20) between the paralogous *ENA* genes in chromosome IV. We detected [*ENA^circles^*] in the young and progeny sub-populations, yet we did not detect the circles in any of the corresponding aged sub-population (Figure 3A, genome coverage plot, PCR and Sanger sequencing), suggesting that [*ENA^circles^*] formed with a high rate and were lost from the population after less than 15-25 cell cycles.

**Figure 3.**
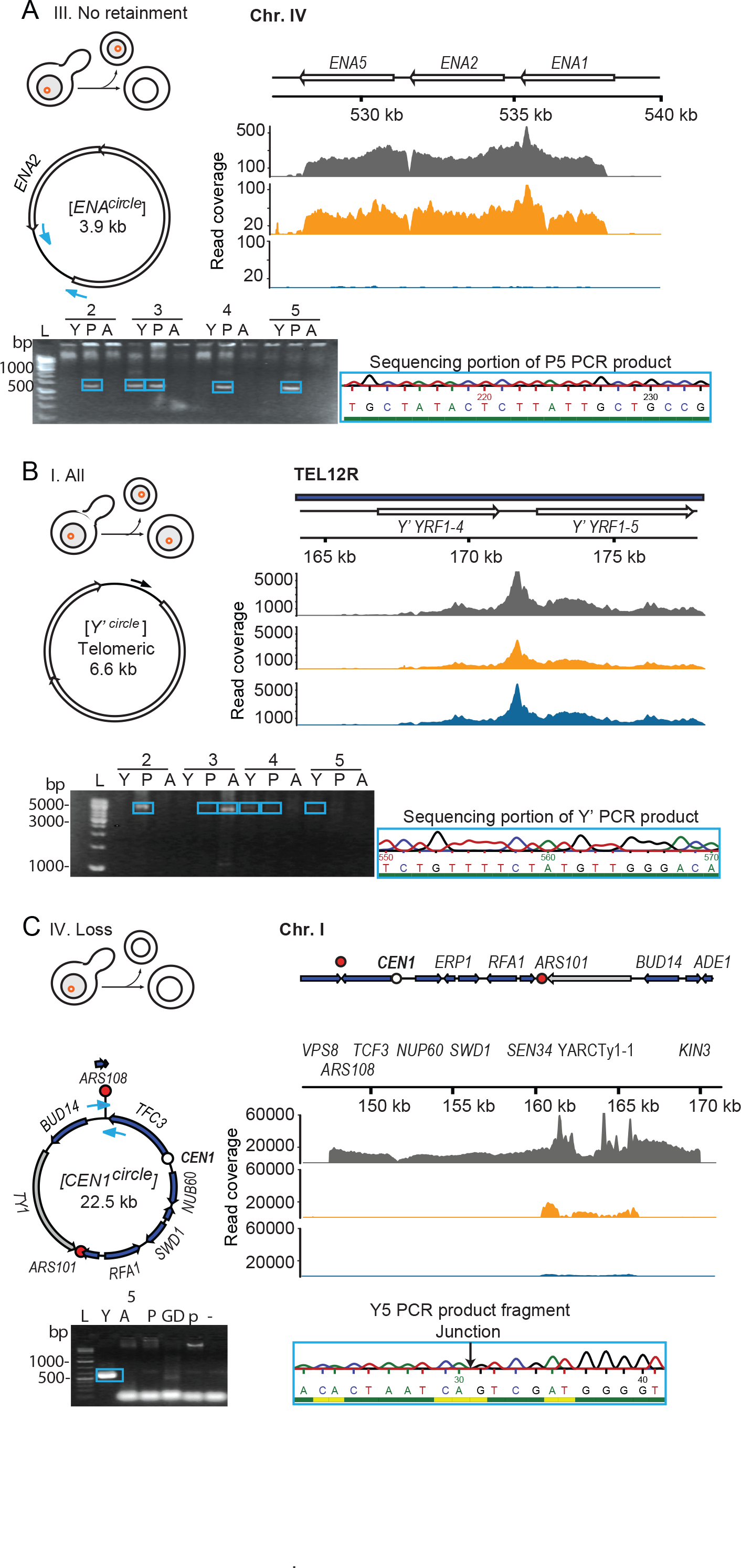
Schematic representation of 3 validated circular DNA with the genomic features (genes shown in blue, repetitive elements shown in white, retrotransposons shown in grey, replication origins shown as red circles and centromere shown as a white circle) together with the read coverage (grey indicates the young population, orange indicates the progeny population and blue indicates the aged population), PCR based gel validation (Y indicates the young population, A indicates the progeny population, GD indicates genomic DNA, P indicates the pUG72 plasmid control, - indicates the negative control and L indicates the ladder) and sanger sequencing. Confirmation of circular DNA in the 2-5 populations using outwards PCR for circular DNA formed from the ENA locus (*A*), the *Y’* telomeric circle (*B*) and a circular DNA spanning the centromere of chromosome 1 (*C*).

Another class of recurring DNA circles without replication origins came from *Y’* telomeric elements (41). Due to the poor alignment of telomeric regions we could not determine the genomic origin of the [*Y’ telomeric^circle^*] based on the short-read sequencing. We therefore used a combination of PCR and Sanger sequencing on purified circular DNA to determine the genomic origin of the [*Y’ telomeric^circle^*] in aged subpopulations. Sanger sequencing revealed that *Y’*-elements could derive from three telomeric (TEL) loci, the right arm of chromosome IV or XV that each carry one copy of the sequence or the right arm of chromosome XII, TEL12R, carrying tandem directed *Y’* repeats. The [*Y’ telomeric^circle^*] sequence matched exactly the middle of the two repeats on TEL12R (Figure 3B, Supplementary File S1). Thus, we anticipate that the [*Y’ telomeric ^circle^*] primarily formed from TEL12R because this circle (without a replication origin) was present in several populations. These traits also suggest high formation rates, which likely is a more frequent event between the Y’-elements in direct repeats on TEL12R than non-homologous ends of *Y’* from other chromosomes. Finally, telomeric *Y’* circles did not carry X’-repeats as expected if they had derived from TEL4R or TEL15R. Taken together, these results suggested that the homologous direct repeat loci TEL12R and *ENA1 ENA2 ENA5* formed circular DNA at high rates.

Circular DNA spanning centromeres are intriguing, as they contain the required genomic signal to segregate faithfully in mitosis and are potential intermediates in the evolution of new chromosomes with altered synteny. We scanned the detected circular DNA (Supplementary Table S2) for the presence of intact centromeres and discovered a 22.5 kb large circular DNA that contained the centromere of chrI, two replication origins, a LTR retrotransposon (Ty1 element) and 7 genes (Figure 3C). Hence, this circular DNA carried all the necessary elements for an equal replication and segregation. Yet, we only found the CEN circle in one young subpopulation and not in the corresponding aged or progeny populations (Figure 3C). This indicates that the [*CEN^circle^*] was lost from the aged yeast populations despite having a centromere. Taken together, these results suggested that the homologous direct repeat loci TEL12R and *ENA1 ENA2 ENA5* formed circular DNA at high rates.

### Replicative aging is associated with loss of genetic heterogeneity from circular DNA

One of the hallmarks of aged yeast cells is the massive accumulation of [*rDNA^circles^*] when compared to their young counterparts (4, 18, 42). To investigate if recurrent non-rDNA circles also accumulated in aging cells, we quantified circular DNA by counting the number of reads aligned to the different circular DNA coordinates and normalizing the read counts, using the median of ratios method (43), as it compensates for the fact that [*rDNA^circles^*] will have a strong influence on the circle specific read counts between the young and aged populations. We then removed circles that only appeared in one yeast sub-population, leading to a set of 13 recurrent circular DNA (Supplementary Table S5). Principal Components Analysis (PCA) revealed that the aged yeast populations clustered tightly together (Figure 4A). Young and progeny sub-populations did not form any apparent clusters in the PCA, supporting that aging sub-populations were less heterogeneous than young. To obtain a better understanding of why old yeast cells became more similar as they aged, we complemented our PCA analysis with a hierarchical clustering of the recurrent circular DNA (Figure 4B). As in the PCA, the sample dendrogram of the hierarchical clustering (Figure 4B, upper dendrogram) showed that aged sub-populations formed a well-defined group, while the young and progeny populations did not form any apparent cluster. When we examined the cells of the hierarchical clustering individually, we noticed an increase in signal for the aged populations in rows representing [*rDNA^circles^*] and [*Y’ telomeric^circles^*] (Figure 4B). The remaining recurrent circular DNA did not show any evidence of accumulation in aged populations, neither for circles containing replication origins nor for those without. Similar results were obtained with another four populations (Supplementary Figure S7A-B).

**Figure 4.**
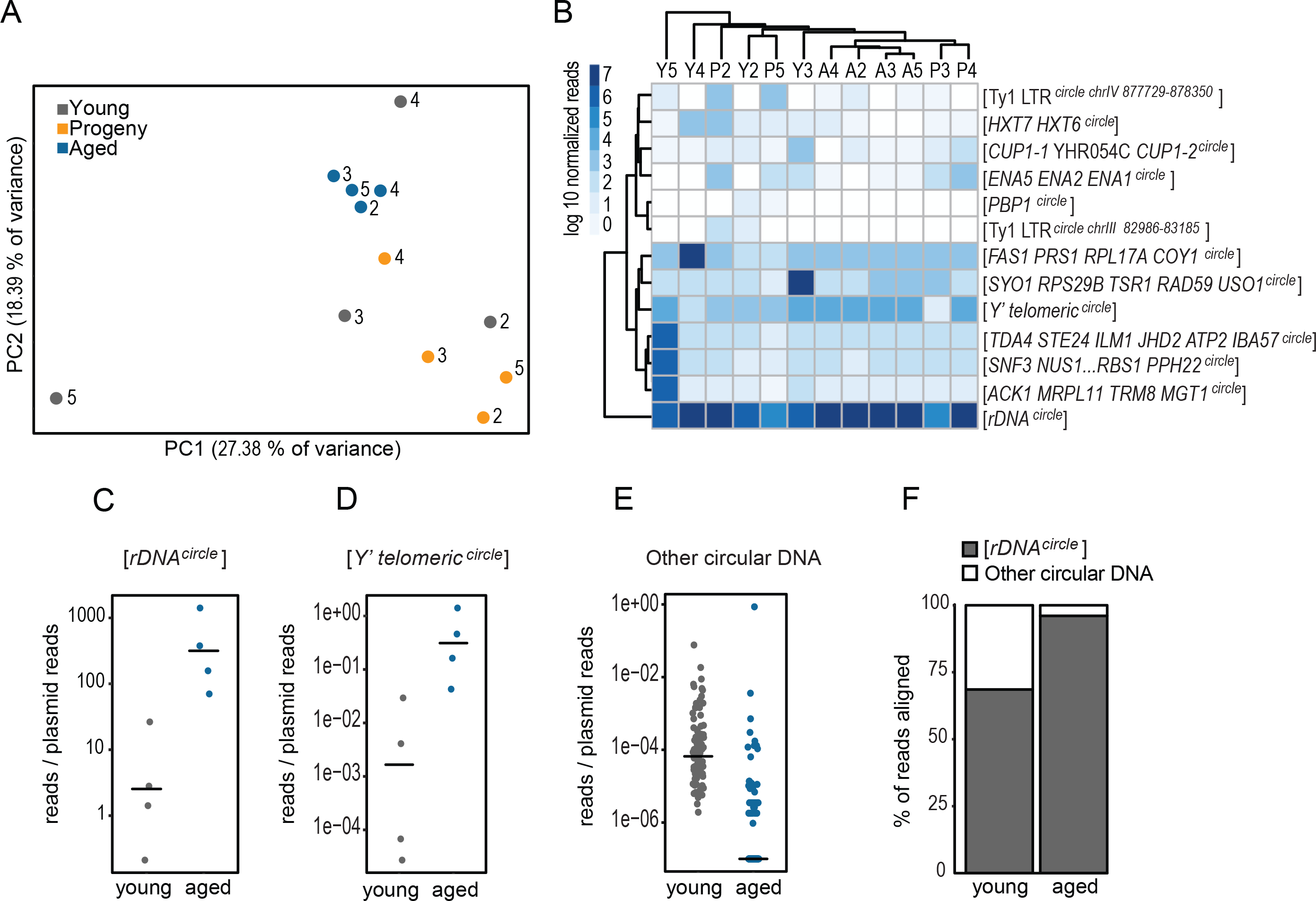
Quantification of circular DNA as cells undergo many divisions. (*A*) Principal component analysis of the circular DNA found in at least two samples of the 2-5 yeast populations. (*B*) Hierarchical clustering of the recurrent circles. Clustering of similar samples is shown in the column dendrogram and clustering of similar circles is shown in the row dendrogram. Read coverage scale is shown in the upper left corner. (*C-E*) Relative levels of [*rDNA^circle^*] (*C*), [*Y’ telomeric ^circle^*] (*D*) and DNA circles from other genomic loci (*E*) calculated by normalizing the read counts to the spike-in plasmids in the 6-10 yeast populations. The horizontal black lines indicate the median value. (*F*) Percentage of reads aligned to the rDNA and to circular DNA from other regions in the 2-5 yeast populations.

To more precisely determine accumulation of [*rDNA^circles^*] and [*Y’ telomeric^circles^*] in the aged samples, we purified and sequenced circular DNA in the presences of spike in plasmids in four populations, samples 6-10. This allowed us to normalize the read count values between the young and aged sub-populations (Figure 4C-E, Supplementary Table S6-8) and make a semi-quantitative measure of circular DNAs between samples. [*rDNA^circles^*] (Figure 4C, Supplementary Table S6) accumulated significantly in the aged samples (median fold-change 432.3, Wilcoxon two-sided test, n=8, *p-value=0.028*). Similarly, we found a significant accumulation of [*Y’ telomeric^circles^*] (Figure 4D, Supplementary Table S7, median fold-change 6427.58, Wilcoxon two-sided test, n=8, *p-value=0.028*), while circular DNA from other parts of the genome did not accumulate significantly (Figure 4E, Supplementary Table S8). Moreover, we used the median of the ratios normalized values to confirm with another normalization method that the majority of the circles lose abundance as cells age (Supplementary Figure S7C). We next addressed how rDNA levels influenced the circular DNA read distribution in the yeast populations (Figure 4F). Young yeast populations were already dominated by reads coming from the rDNA locus (median percentage of aligned reads 67.57%, range: 47.78-93.23%). Still, the percentage increased drastically on the aged yeast populations (median 98.55%, range: 87.37-99.53%), where the reads originating from [*rDNA^circles^*] dominated the samples over circular DNA formed from other genomic loci. Finally, we confirmed the accumulation of [ *rDNA^circles^*] on the 2-5 aged populations using qPCR (Supplementary Figure S8).

To confirm that DNA circles from other parts of the genome did not accumulate in aged cells, we measured the level of purified circular DNA upon exonuclease treatment with qPCR. Quantification of two circles that did not accumulate in populations 2-5, [*CUP1/2^circle^*] and [*GAP1^circle^*], from two independent MEP enriched yeast populations confirmed that [*CUP1/2^circle^*] and [*GAP1^circle^*] levels either stayed unaltered or decreased in aged subpopulations compared to young subpopulations (Supplementary Figure S9A-C). Taken together, these results show that aging cells primarily accumulated [*rDNA^circle^*] and that the number and level of non-rDNA circles generally decreased as cells aged.

### The [*HXT6/7^circle^*] alter inherence pattern when selected for and is detected prior stable chromosomal *HXT6/7* amplifications

Circular DNAs are known to enrich under conditions where they are selected for (13, 44). We speculated that circular DNA could change inherence pattern under different selective regimes from failing to be maintained (class III and IV) towards occurring in all cell types (class I), and vice versa. To study the inheritance pattern of a circular DNA under selective pressure, we followed the distribution of [*HXT6/7^circle^*] in a population of cells exposed to prolonged glucose limitation. The *HXT6* and *HXT7* genes encode high affinity glucose transporters, and growth under glucose limitation is known to select for chromosomal amplification of the *HXT* genes in the *HXT6 HXT7* locus to a *HXT6 HXT6/7 HXT7* locus (39, 45, 46). The *HXT* locus can also excise as a [*HXT6/7^circle^*] with the 5’-end of *HXT6* and the 3’-end of *HXT7* on it (Figure 5A, Supplementary Table S2) (20, 28), but the [*HXT6/7^circle^*] has so far not been associated to amplification of the *HXT6 HXT7* locus.

**Figure 5.**
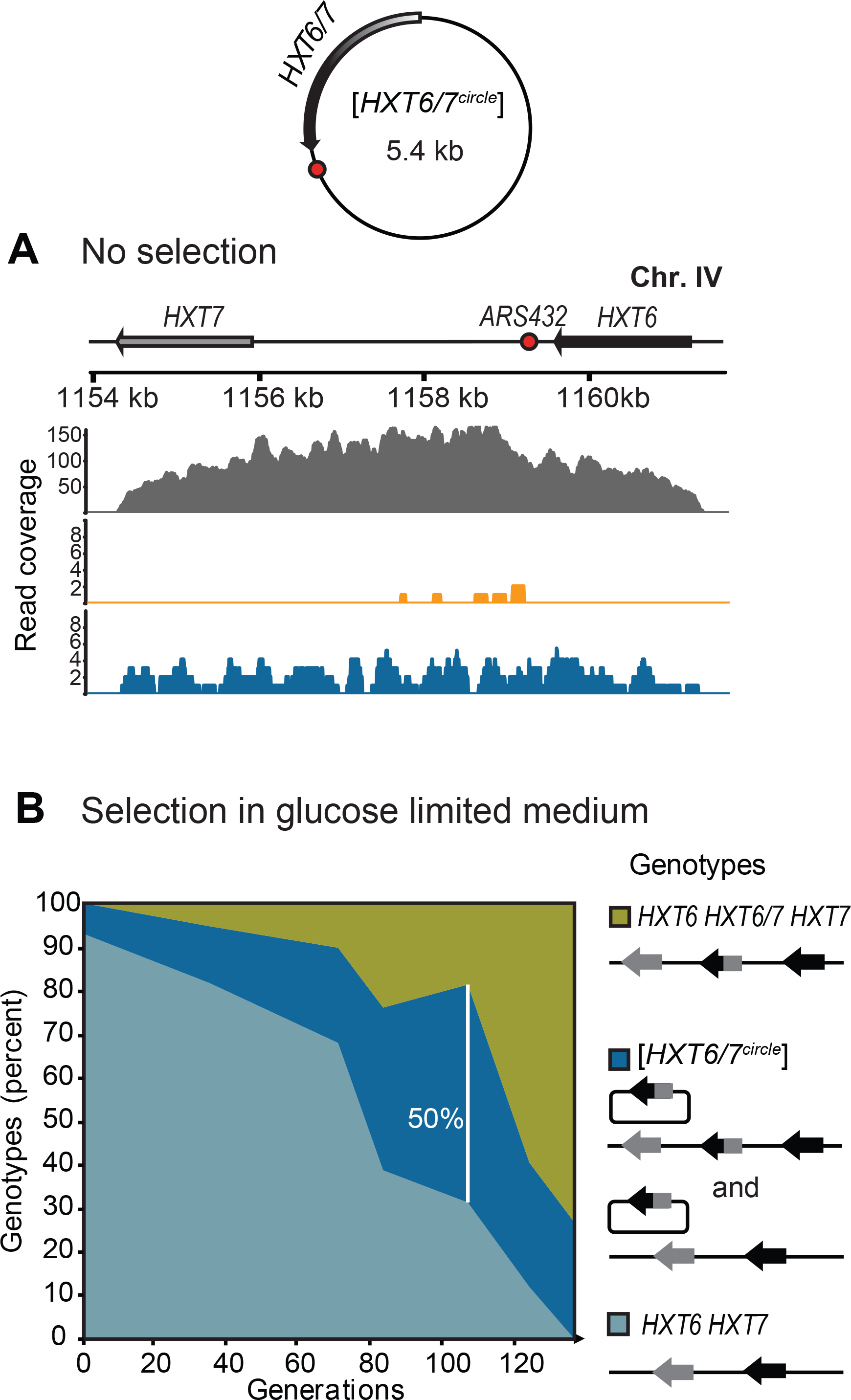
Segregation behaviour of circular DNA formed from the *HXT6/7* locus under non-selective and selective pressures. (*A*) Read coverage plot of the [*HXT6/7^circle^*] under no selection (detected in sample 4). The read coverage is shown for the young (grey), progeny (orange) and aged (blue) sub-populations. The replication origin is shown as a red circle in the schematic diagrams. The paralogous genes *HXT6* and *HXT7* are shown in grey and black respectively. (*B*) Different genotypes of the *HXT6/7* (wild-type shown in light blue, circular amplification shown in dark blue and chromosomal amplification shown in green) under glucose limitation during 135 generation (genotyped at generations: 0, 35, 71, 82, 106, 124 and 135).

The [*HXT6/7^circle^*] was discovered four subpopulations, but only one time in the progeny population, suggesting that [*HXT6/7^circle^*] had a segregation bias towards the mother cells as other class II circles (Figure 5A). We next investigated if the [*HXT6/7^circle^*] would be inherited to daughter cells if these were under selection for keeping the [*HXT6/7^circle^*]. This was done by tracking the *HXT6 HXT7* genotypes of individual cells during 135 generations of glucose limitation (Figure 5B) at various time points (0, 35, 71, 82, 106, 124 and 135). Initially, the majority of the genotypes were wild-type (93.02% wild-type), with a minor fraction of cells carrying the [*HXT6/7^circle^*] (6.98% circle carriers) and no stable chromosomal amplifications. As selection progressed, the genotypes switched towards cells carrying the [*HXT6/7^circle^*] and eventually shifted to cells with chromosomal *HXT6 HXT6/7 HXT7* amplifications dominating the population. The fact that a large number of cells (50% of the cells at generation 106) were found to contain the [*HXT6/7^circle^*] suggested that the [*HXT6/7^circle^*] was inherited by daughter cells under selective pressure. Exponentially growing cells consist of 50% young daughter cells, which have never divided, 25% that have been mothers once, 12.5% have been mother twice and so forth. Therefore a large proportion of the cells carrying [*HXT6/7^circle^*] under glucose limited conditions must be daughters that have received the [*HXT6/7^circle^*] from their mother. In short, inherence patterns of circular DNA can shift under different selective regimes. Furthermore, the progressive accumulation of the [*HXT6/7^circle^*] phenotype, followed by stable amplification genotype suggested that circular DNA serve as intermediate units for the formation of stable chromosomal amplifications.

## DISCUSSION

Our study suggests that circular DNA of chromosomal origin can follow three different inherence patterns in cells as they undergo replication. A majority of circular DNA is lost and only a small fraction of all circular DNA appears to be maintained as cells replicate. A subgroup of the latter are [rDNA*^circles^*] that accumulate over other circular DNA species as cells age. This eventually results in aged cells with a relatively low genetic variation from circular DNA and strong accumulation of [rDNA*^circles^*].

Circles found in both young and aged cells are characterized by replication origins sugges ting that replication is essential for maintenance of circular DNA. This finding supports previous studies which have used acentric plasmids containing replication origins as a model to study the behaviour of circular DNA during cell division (8, 11). When cells containing these plasmids were growth on non-selective medium, there was a segregation bias towards the mother cell, leaving the daughter cell with no copies of the plasmid (8, 11).

Our results suggest that the majority of replicating circles do not accumulate to the same extent as [rDNA*^circles^*] and [*Y’* telomeric*^circles^*] in aging cells. We speculate that the discrepancy between [rDNA*^circles^*] and other circles can be caused either by an accumulation mechanism that is specific for [rDNA*^circles^*] or a very high formation rate from the rDNA locus. Barral and co-workers have suggested an active retention of [*rDNA^circles^*] and other circular DNA in mother cells through the binding of the circular DNA to the SAGA complex and the nuclear pore complexes (8). The model does not distinguish between [rDNA*^circles^*] and other circular DNA with replication origins and can therefore not explain their discrepancy. Circular rDNA formation has been shown to increase as cells age (5, 18) and could thus contribute to their accumulation. The rDNA locus is sequestered in the nucleolus in another enzymatic environment than other parts of the nucleus, with other DNA polymerases that can also influence formation rate (9). It seems feasible that the physical separation of nucleus and nucleolus will also allow for a separate segregation mechanisms as recently suggested (5). Recent studies indicate that the level of [rDNA*^circles^*] also depends on the length of the chromosomal rDNA repeat (9). We found that [rDNA*^circles^*] are already present in young cell populations, presumably carried by a minor fraction of aged cells. Then, as the cells age the [rDNA *^circles^*] become the dominant circular DNA form in the population. The accumulation of rDNA circles in aging cells over other circular DNA might partly be a consequence of this regulation of rDNA copy number but how the self-limiting production of [rDNA*^circles^*] interplays with accumulation in aging cells is so far unknown. Hence, we can currently not distinguish i f the high level of [rDNA *^circles^*] over other maintained circles (Figure 4F) is caused by different segregation mechanisms (passive or active retention) or because they have different formation rates but the same segregation mechanism.

An important exception to the inherence patterns found for most non-rDNA circles, are circles from genes in direct repeats (*ENA1 ENA2 ENA5, Y’* telomeric and *HXT6 HXT7*). Neither *ENA1 ENA2 ENA5* or *Y’* telomeric circles carry replication origins and the fact that these circles are found recurrently in independent populations (Figure 3); suggest that they are formed *de novo* with high rates. *CUP1 CUP2* and *HXT6 HXT7* carry replication origins, and are also found recurrently, suggesting that circles from direct repeats in general have a high formation rate. Hence, our data supports the notion (4, 40) that three factors determine how a circular DNA is maintained in a cell lineage: the rate by which it is formed from its chromosomal locus, its ability to replicate and its mode of segregation, as recently.

Interestingly, we find that aged cells have a lower number of different circular DNA than young cells. This suggests that yeast cells lose circular DNA genetic heterogeneity through aging and that the genetic variation upon which selection can act becomes lower with age. Young cells have a larger pool of genetic variation than old do, thereby theoretically allowing young cells to adapt and respond a wider set of condition than old cells. Furthermore, genes linked to replication origins and direct repeats are more likely to reside on circular DNA, evade the 1:1 segregation pattern, and thereby contribute to genetic variation in the form of circular DNA. This implies that replication origin linked genes might evolve faster, as they exist longer in a given populations when not under selective pressure. Several of the genes in direct repeats are known to evolve fast in yeast populations (47) and circular DNA could be an intermediate between deletions and amplifications. While we cannot rule out the possibility that we did not detect a minor fraction of the circular DNA found in the aged populations, our computational experiments suggest that we found the majority of the circular DNA present in the young, progeny and aged populations already at a 20% of the sequencing depth. Hence, we are confident that our circular DNA loss rate is not overestimated. An example of circles with high formation rates and replication origin is the [*HXT6/7^circle^*] that carries a hybrid of two high-affinity hexose transporter genes from the *HXT6 HXT7* locus. We have found this circle recurrently in our previous whole genome circles studies (20, 28) and the circle appears repeatedly in independent populations. Results from others have shown that glucose limitation selects for amplification of the *HXT6 HXT7* locus to *HXT6 HXT6*/*7 HXT7* (45, 46). Our data suggest that under selective pressure the [*HXT6/7^circle^*] segregation patterns change and the proportion of cells with [*HXT6/7^circle^*] increase in the population until cells with stable chromosomal amplifications arise and take over the population. We propose that the [*HXT6/7^circle^*] can be an intermediate in the formation of the more stable *HXT6 HXT6/7 HXT7* chromosomal amplification, and that circular DNAs can facilitate a fast adaptation to selective conditions via a high formation rate, replication and mis-segregation.

Amplification of genes in direct repeat through a circular intermediate might also apply to other genes in direct repeat in both yeast and other eukaryotes such as human, where genes in direct repeat are known to amplify and lead to disorders such as Charcot-Marie-Tooth or the Williams-Beuren syndrome (48). In yeast, the [*Y’* telomeric*^circle^*] has been suggested to be the intermediate in telomeric extensions in type I survivors of telomerase deficient cells (41). However, the origin of [*Y’* telomeric*^circles^*] has been unknown but our study suggests they arise primarily from the *Y’*-repeat in TEL12R, possibly due to the structure with two tandem *Y’* repeats.

We have not addressed the fate of class IV circles, which were only detected in young cells. Most of these circles might not have replicated due to the lack of origins hence only existed in the one copy recorded in the young population. Alternatively, class IV circles might have replicated rarely and could have diffused to daughter cells. Because we only recorded circular DNA from 3% of the progeny populations, our analysis will have missed these circles in the progeny. We also found that a [*CEN1^circle^*] was lost during replicative aging. While this circle was expected to segregate 1:1, cells carrying the [*CEN1^circle^*] might have been lost from the aged population due to decrease in fitness of cells carrying them. Cells with a [*CEN1^circle^*] are expected to have lost major fragments of chromosome I, which is expected to result in a much lower fitness compared to wildtype diploids.

In conclusion, our whole genome survey of circular DNA in *S. cerevisiae* supports three factors determine how a circular DNA is maintained in a cell lineage: the rate by which it is formed from its chromosomal locus, its ability to replicate and its mode of segregation (4). Our data suggest that rDNA circles are the dominating circles in replicative aged cells and that young cells have the largest variety of different circular DNA. There appears to be large differences in the propensity of different parts of the genome to evolve through circular DNA, since loci linked to direct repeats and replication origins are more likely to reside on circular DNA. Selection can furthermore change the inherence pattern of a circle and lead to rapid propagation of circle-carrying cells in a population. These observations are not only relevant for yeast but can also be important for our understanding of aging in other organisms and our insight into evolution of cancer cells through circularized proto-oncogenes.

## AVAILABILITY

Circle-Map Repeats is open source and available in the GitHub repository (https://github.com/iprada/Circle-Map)

## ACCESSION NUMBERS

The raw sequence data have been deposited on the Sequence Read Archive and can be accessed with no restrictions using the BioProject ID PRJNA593745.

## SUPPLEMENTARY DATA

Supplementary Data are available at NAR online.

## AUTHOR CONTRIBUTIONS

H.D.M, J.H and B.R designed the study. H.D.M, C.E.L and S.A performed the laboratory experiments. I.P.L and R.A.H performed the computational experiments with assistance from L.M. I.P.L and B.R analyzed the data. I.P.L and B.R wrote the manuscript with input from all the authors.

## ACKNOWLEDGEMENTS

We thank Maitreya Dunham for the CEN.PK G6 population propagated under glucose limitation. This research, HDM and BR were supported by a research grant (VKR023513) from the VILLUM Foundation. IPL and BR were supported by (FNU 6108-00171B). We thank GenomeDK and Aarhus University for computational resources.

---

**Supplementary figure S1.**
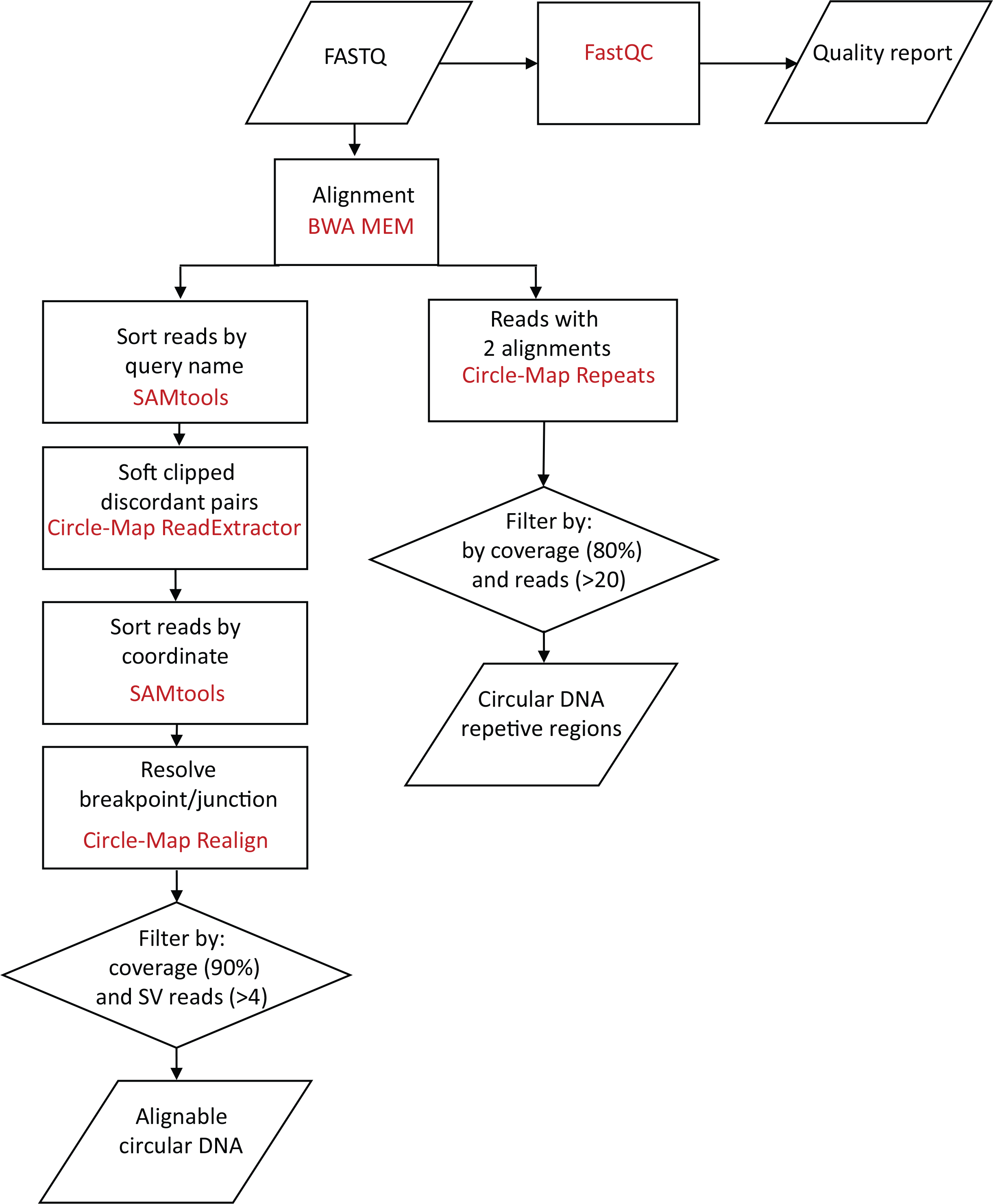
Flowchart describing the bioinformatics workflow conducted to obtain circular DNA coordinate files from the raw data. Parallelograms indicate input/output files. Squares indicate programs used during the workflow. Rhombuses indicate filters applied to the data.

**Supplementary figure S2.**
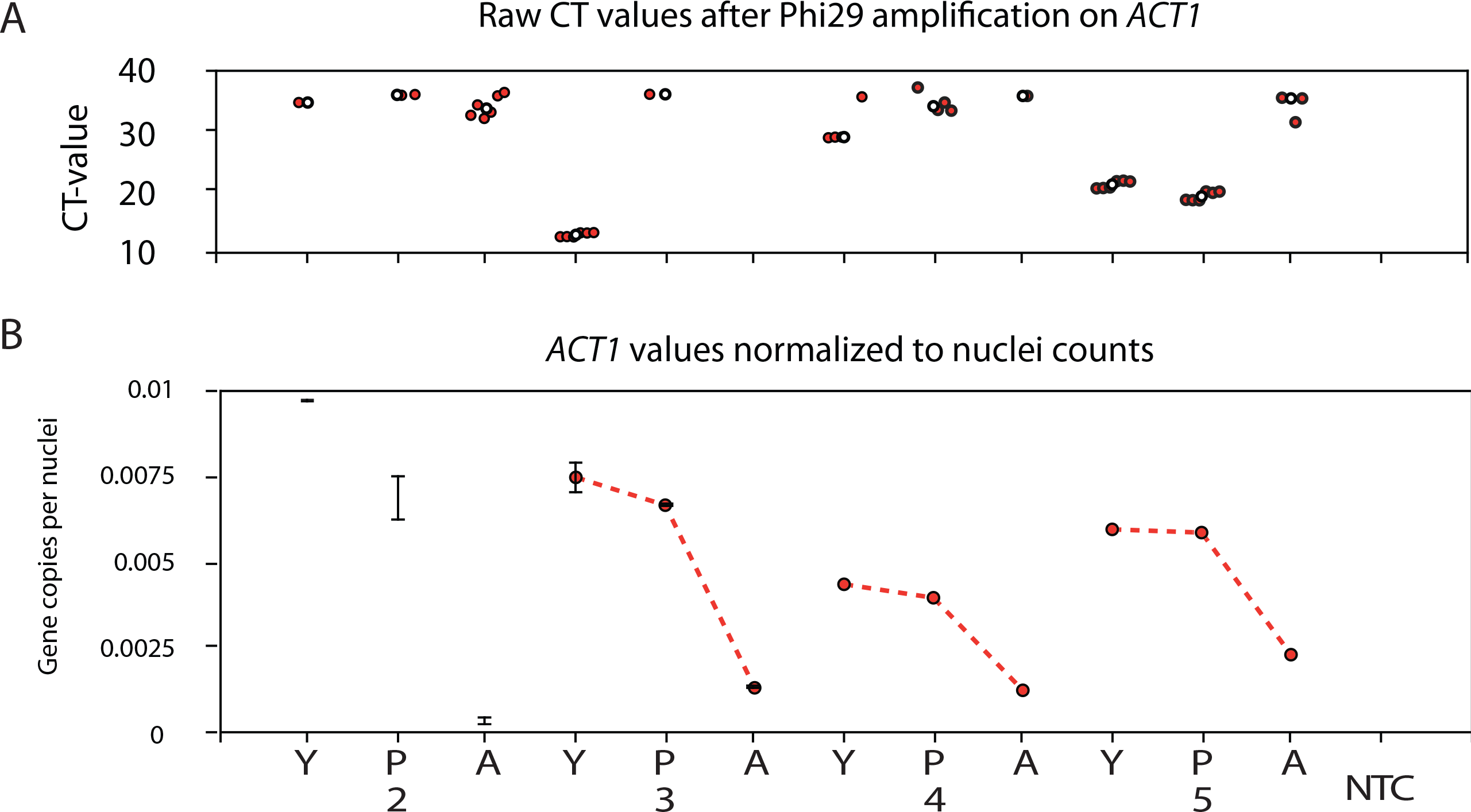
(*A*) Confirmation of linear DNA removal with qPCR on the dosage sensitive gene *ACT1.* The upper panel shows the raw cycle threshold values (CT) for *ACT1* qPCR. (*B*) *ACT1* values normalized to the nuclei count forthe B2-5 samples. Young samples are indicated as Y, progeny samples are indicated as P and aged samples are indicated as A. NTC indicates non template control.

**Supplementary figure S3.**
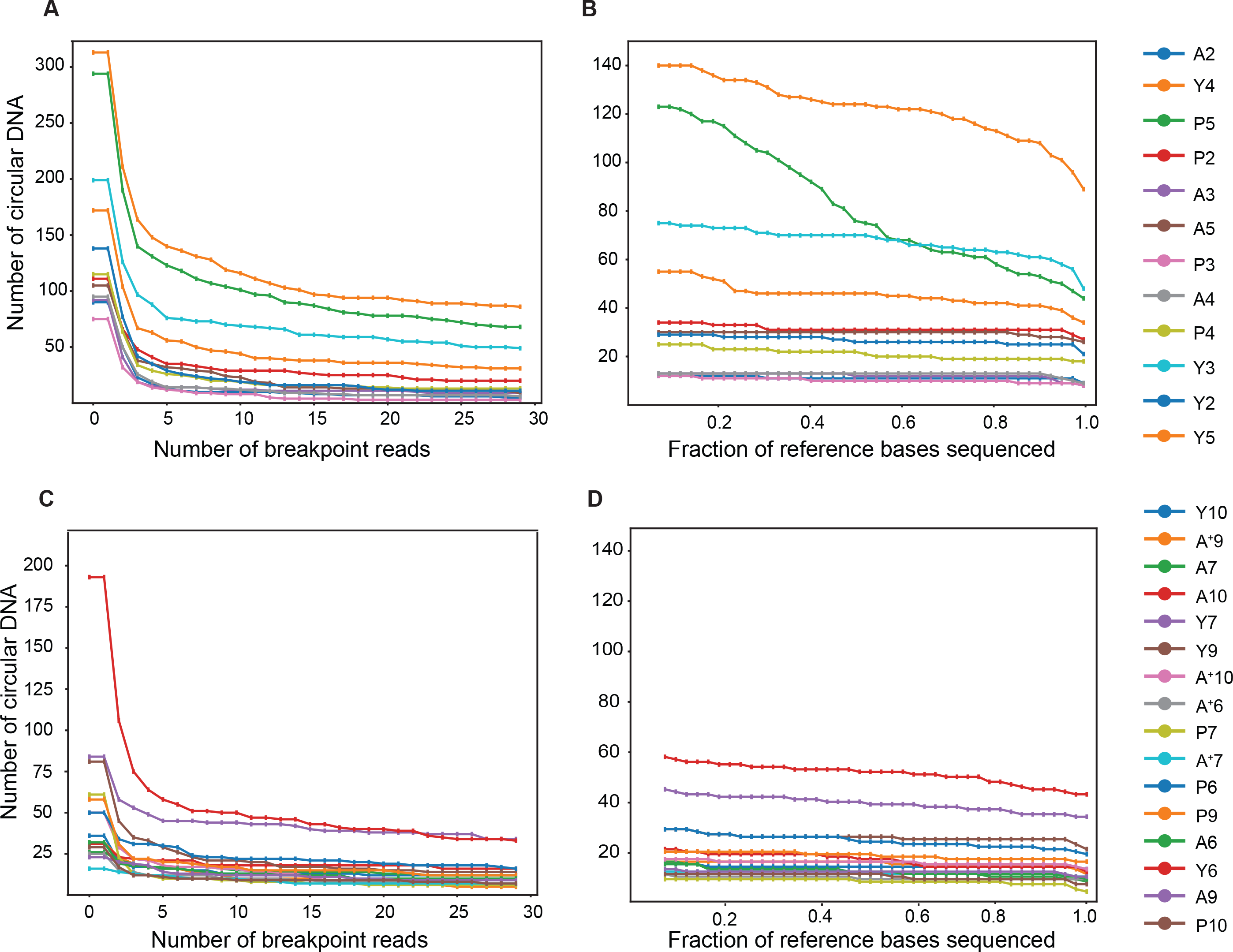
Effect of breakpoint reads and circular DNA sequencing coverage on circular DNA detection. Young populations are indicated with Y, progeny populations are indicated with P, aged populations are indicated with A and aged populations selected with the biotinylation program are indicated with A^+^. (*A*) Numberof detected circular DNA against the number reads crossing the circular DNA breakpoint (split and discordant reads) in the 2-5 yeast populations. (*B*) Numberof detected circular DNA against the sequencing coverage within the detected circular DNA coordinates in the 2-5 yeast populations. (*C*) Numberof detected circular DNA against the numberof reads crossing the circular DNA breakpoint (split and discordant reads) in the 6-10 populations. (*D*) Numberof detected circular DNA against the sequencing coverage within the detected circular DNA coordinates in the 6-10 yeast populations.

**Supplementary figure S4.**
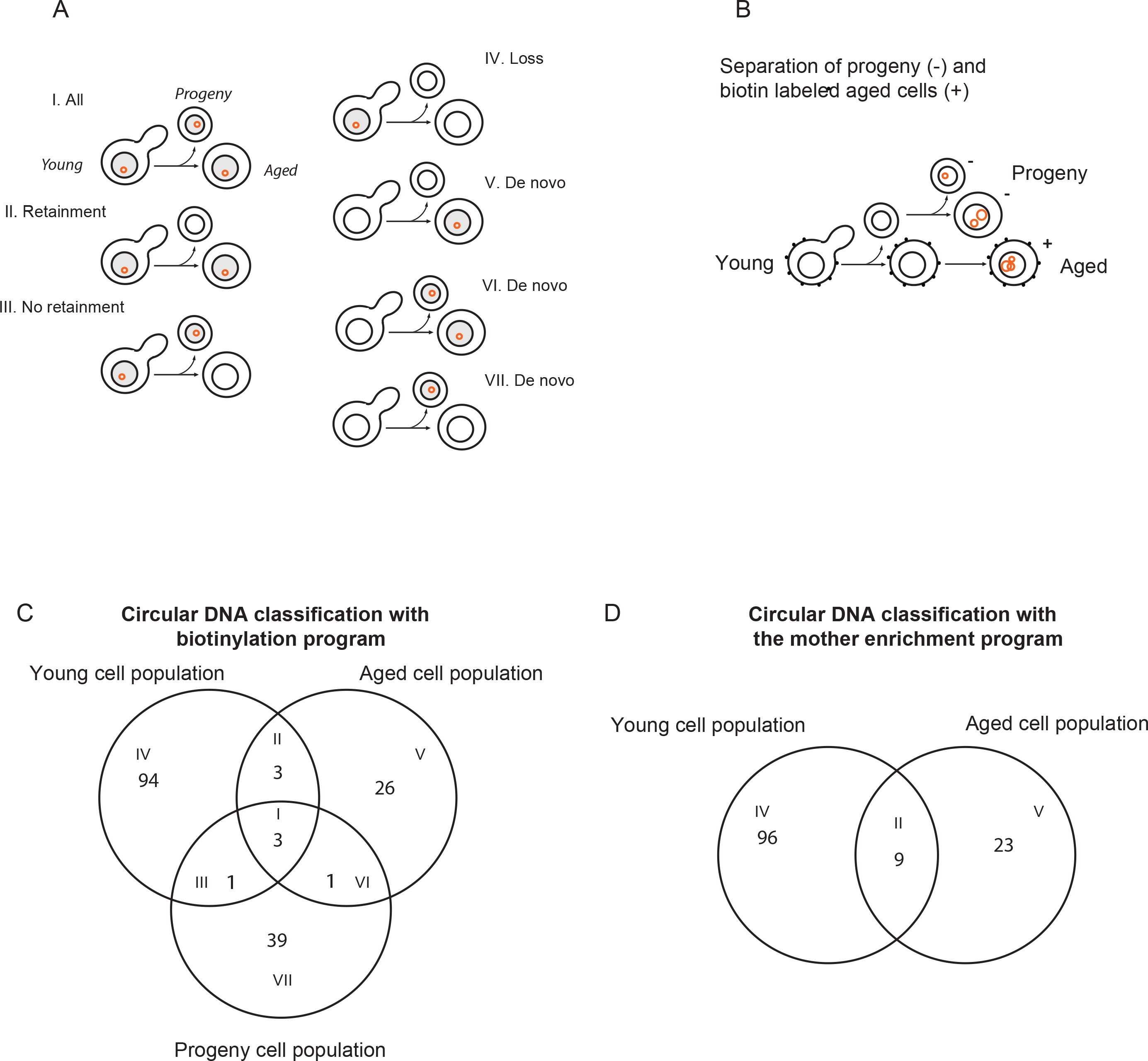
Experimental design and circular DNA segregation patterns forthe 6-10 yeast populations. (*A*) Schematic representation of the different circular DNA segregation patterns. (*B*) Schematic overview of young, progeny and aged cell separation from 4 yeast populations using the biotinylation program. (*C*) Venn diagram displaying the different circular DNA segregation patterns found on the 6-10 yeast populations separated with the biotinylation experiment. (*D*) Venn diagram displaying the different circular DNA segregation patterns found in the 6-10 populations separated using the mother enrichment program.

**Supplementary figure S5.**
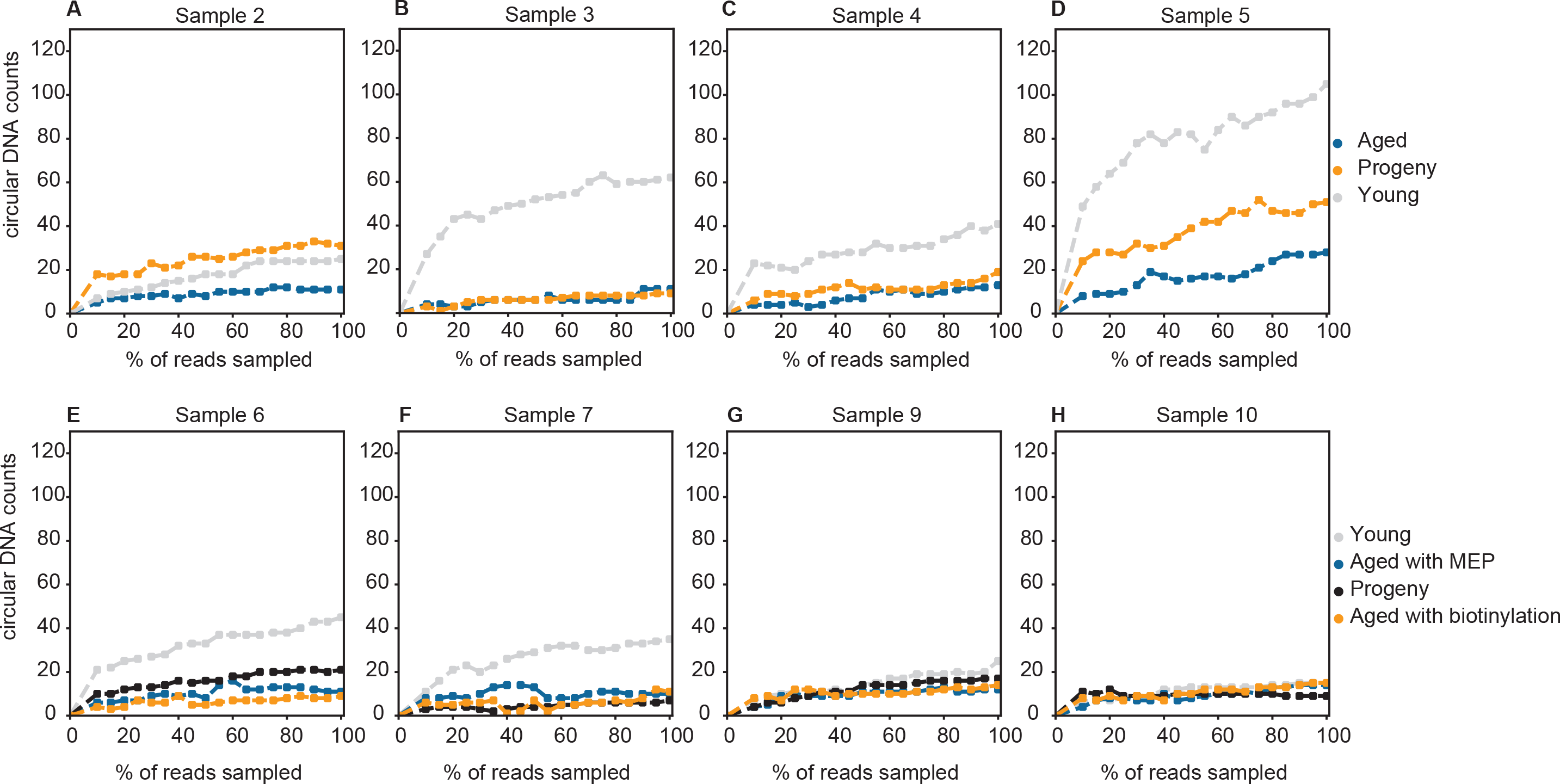
Effect of sequencing depth on the numberof circular DNA detections. (*A-D*) Numberof detected circular DNA against the percentage of reads sampled from the alignment file in the 2-5 yeast populations. (*E-H*) Numberof detected circular DNA against the percentage of reads sampled from the alignment file in the 6-10 yeast populations.

**Supplementary figure S6.**
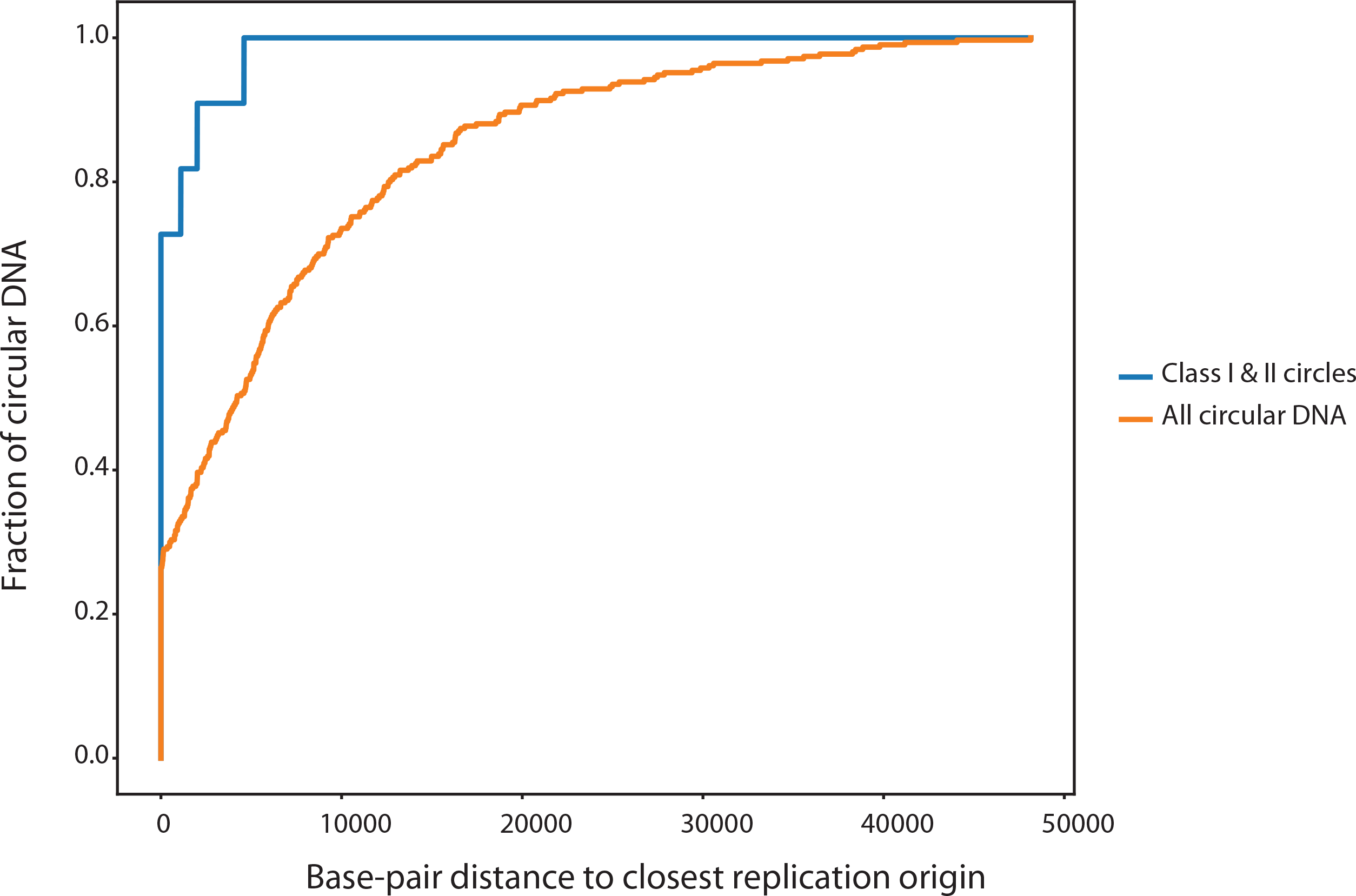
Empirical cumulative distribution functions of the base-pair distance between every circular DNA and the closest replication origin. The class I & II circles are indicated in blue and all the DNA circles are indicated in orange.

**Supplementary figure S7.**
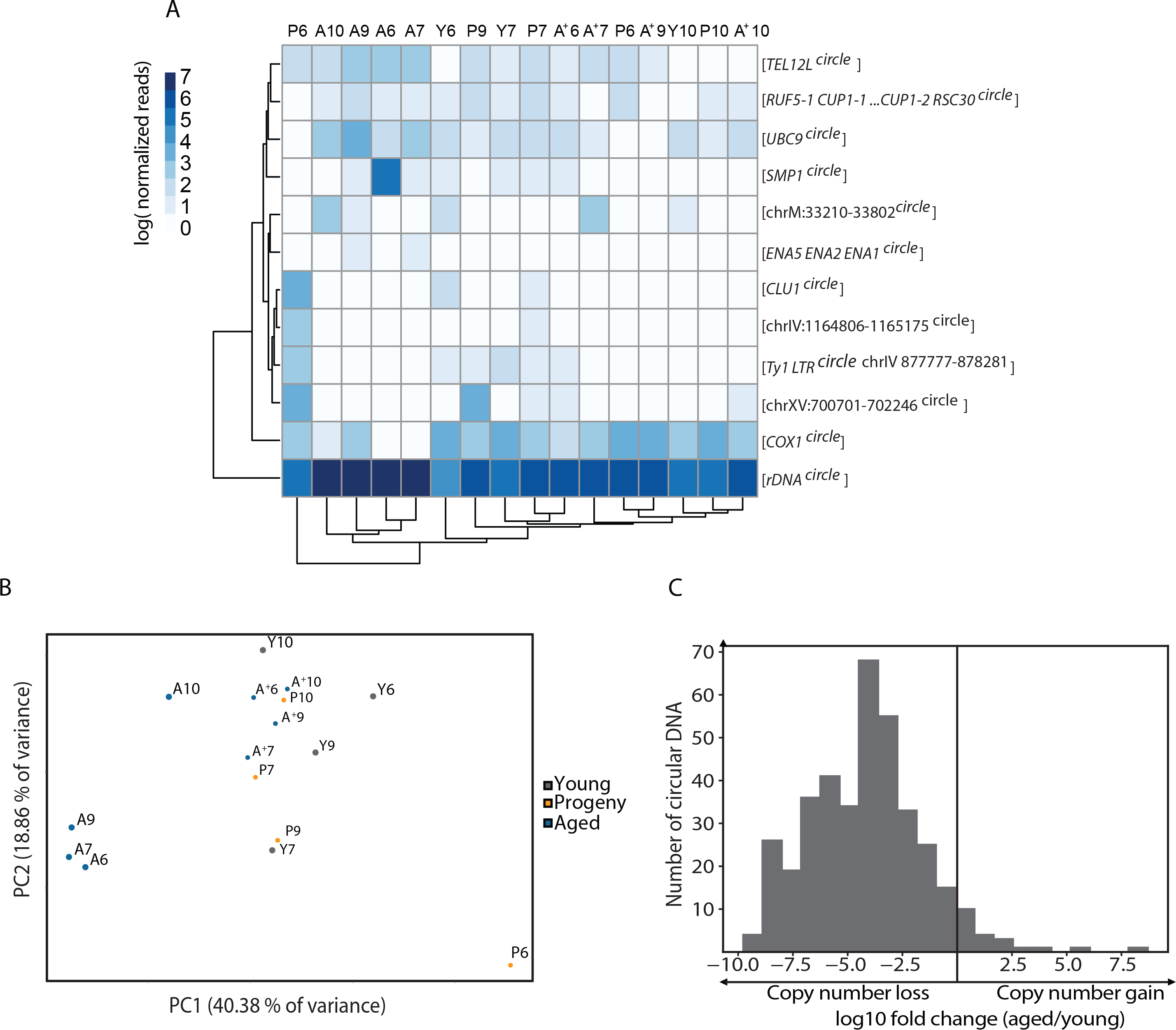
Quantification of circular DNA as cells undergoes many divisions in the 6-10 populations. (*A*) Hierarchical clustering of the circular DNA detected in at least two samples of the 6-10 populations. The left dendrogram shows the clustering order of the circles (rows), with circle names indicated on the left. The dendrogram belowshows the clustering orderof the samples (column), with the sample names indicated in the top part of the plot (Y: young, P: progeny; A: aged with MEP and A^+^: aged with biotynilation). The normalized read coverage is shown on the upperleft part. (*B*) Principal component analysis of the normalized read values of the circular DNA detected in at least two samples of the 6-10 yeast populations. (*C*) *Log* Fold-change distribution between young and aged samples forall the circular DNA detected in all the yeast populations 2-10.

**Supplementary figure S8.**
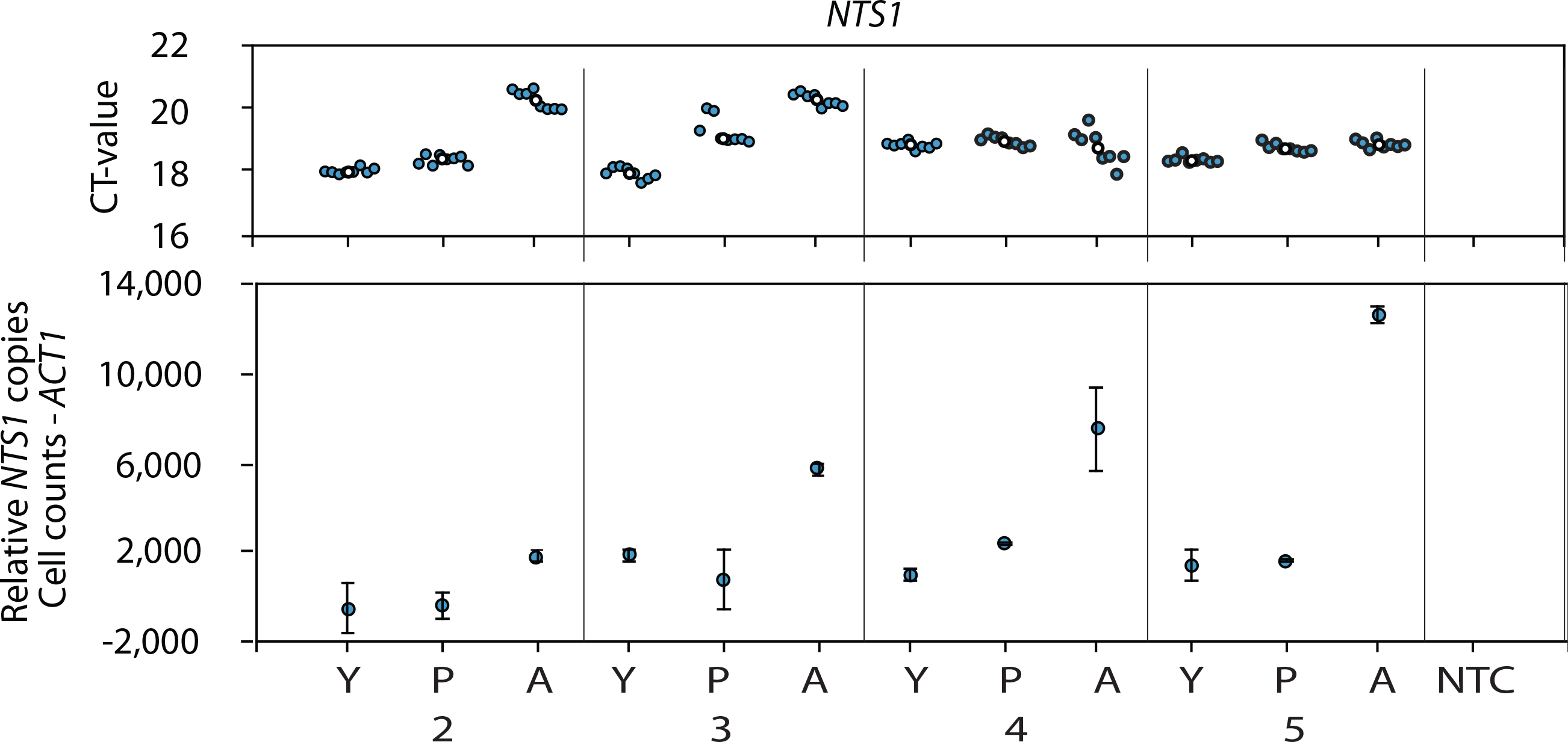
Quantification of [*rDNA^circle^*] levels on the young (Y), progeny (P) and aged (A) subpopulations of the 2-5 samples by performing qPCR on the *NTS1* gene after removal of the linear DNA. The upper panel shows the raw cycle threshold values (CT) and the lower panel shows the *NTS1* values normalized to the cell counts and the dosage sensitive gene *ACT1*. NTC indicates non template control.

**Supplementary figure S9.**
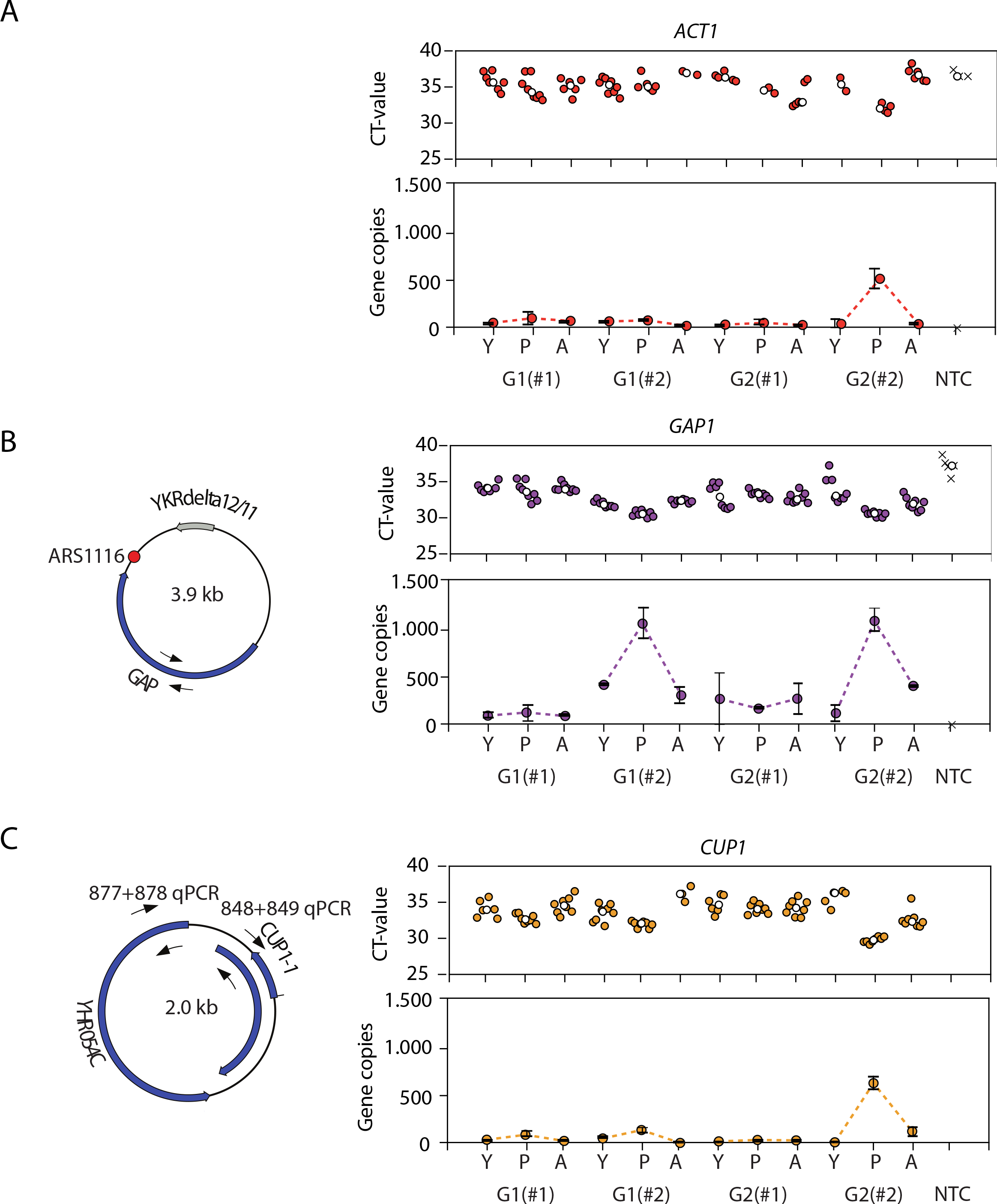
Quantification of the *GAP1* and *CUP1* levels on the young (Y), progeny (P) and aged (A) subpopulations of the G1-G2 after removal of the linear DNA. (*A*) Confirmation of the linear DNA removal with qPCR on the dosage sensitive gene *ACT1*. The upper panel shows the raw cycle threshold values (CT) and the lower panel shows *ACT1* copies normalized to the cell counts. (*B*) Schematic representation of the genomicfeatures present on the [*GAP1^circle^*] (right part). The black arrows indicate the primerlocation used forthe qPCR. The right part shows the qPCR CT values (upper panel) and the cell count normalized gene copies (lower panel). (*C*) Schematicrepresentation of the genomicfeatures present on the [*CUP1^circle^*] (right). The black arrows indicate the location of the primerpairs used forqPCR quantification. The right part shows the qPCR CT values (upper panel) and cell count normalized gene copies (lower panel).The white dots in the upper panels show the median value. The # in the labels of the axis shows technical replicates.

**Supplementary Table S1.**
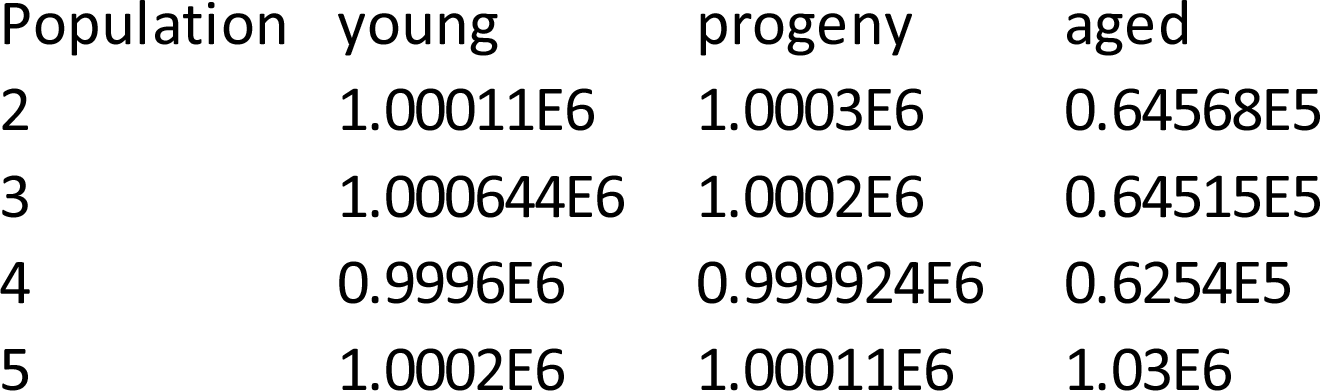
Tab separated file containing the cell counts indicating the numberof cells recovered from every young, progeny and aged population. Ordered from left to right, the columns contain the following information: 1, yeast population; 2, cell counts in the young subpopulation; 3, cell counts in the progeny subpopulation and 4, cell counts in the aged subpopulation.

**Supplementary Table S2:**
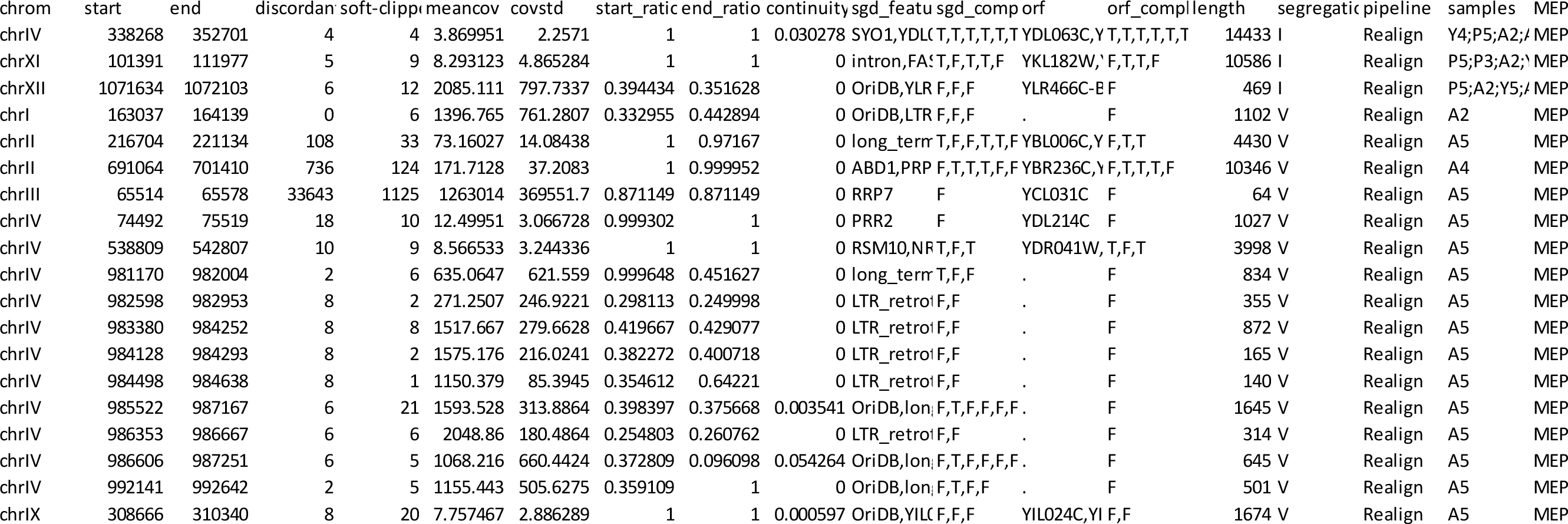
List of all circular DNA recorded in the 8 yeast populations. Top 20 lines shown. Tab separated file containing the chromosomal coordinates, coveragestatistics and genomicfeatures of the detected circular DNA. Ordered from left to right, the columns contain the following information: 1,chromosome; 2, start coordinate; 3, end coordinate; 4, numberof discordant read pairs; 5, numberof split reads; 6, mean coverage within the detected coordinates; 7, standard deviation of the coverage within the detected coordinates; 8, read cove rage ratio at the start coordinate; 9, read coverage ratio at the end coordinates; 10, fraction of reference bases not covered by sequencing reads within the detection coordinates; 11, genomicfeatures contained on the circle; 12, True (indicated as T) and False ( indicated as F) values indi cating whetherthe genomic feature overlaps the circle at 100%; 13, ORFs contained on the circle; 14, True ( indicated as T) and False ( indicated as F) values indicating whether the ORFs overlaps the circle at 100%; 15, length of the circle; 16, classification of the segregation mechanism during cell division; 17, Circle-Map algorithm (Realign or Repeats) used for detection of the circle; 18, samples number identifiers(2–10) and subpopulations (A: aged, P: progeny, Y: young) were the circle was detected and 19, purification experiment were the circle was detected ( MEP: Mother Enrichment Program, Biotin: Biotinylation program and Rep-MAP: replicated Mother Enrichment Program).

**Supplementary Table S3:** Top 20 lines shown. Tab separated file containing the *cis* acting element binding sites tested for overrepresentation between circular DNA classified as lost and circular DNA classified as retained (class I and II). Ordered from left to right, the columns contain the following information: 1, *cis* acting element name tested; 2, regularexpression used forsearching the *cis* acting element in the reference sequence; 3, Fisher’s exact test odds ratio; 4, Fisher’s exact test *p-value*; 5, adjusted *p-value*; 6, numberof circular DNA containing the *cis* acting element in the retained group; 7, numberof circular DNA not containing the *cis* acting element in the retained group; 8, numberof circular DNA containing the *cis* acting element in the lost group and 9, numberof circular DNA not containing the *cis* acting element in the lost group.

| Gene | Motif_reg | OddsRatio | P-value | P-Adjust | In_Retained | Not_in_Retained | In_lost | Not_in_lost |
| --- | --- | --- | --- | --- | --- | --- | --- | --- |
| ABF1 | [AG]TC[AC | 1.5 | 0.5319 | 1 | 5 | 6 | 70 | 126 |
| ACE2 | ACCAGC | 1.6337 | 0.5355 | 1 | 6 | 5 | 83 | 113 |
| ADR1 | G[AG]GG | 0.4 | 0.2458 | 1 | 9 | 2 | 180 | 16 |
| AFT1 | [TC][AG]C | 1.9219 | 0.4017 | 1 | 3 | 8 | 32 | 164 |
| ARO80 | CGG[GATC | 2.1595 | 0.3816 | 1 | 3 | 8 | 29 | 167 |
| ASH1 | [AT][AT][C | 1.0402 | 1 | 1 | 8 | 3 | 141 | 55 |
| AZF1 | AAGAAAA | 1.6795 | 0.514 | 1 | 5 | 6 | 65 | 131 |
| BAS1 | TGACTC | 1.6414 | 0.5166 | 1 | 5 | 6 | 66 | 130 |
| YAP1 | TTAGT[CA | 1.3269 | 0.7387 | 1 | 4 | 7 | 59 | 137 |
| CAT8 | CC[AG][G/ | 0.9145 | 1 | 1 | 3 | 8 | 57 | 139 |
| CBF1 | TCACGTG | 0.88 | 1 | 1 | 1 | 10 | 20 | 176 |
| CHA4 | CGG[GATC | 2.0332 | 0.2753 | 1 | 4 | 7 | 43 | 153 |
| CRZ1 | G[GATC]G | 1.52 | 0.6397 | 1 | 2 | 9 | 25 | 171 |
| CUP2 | [ACT]T[AC | 2.5083 | 0.2214 | 1 | 8 | 3 | 101 | 95 |
| GATA | GATAAG | 1.0019 | 1 | 1 | 5 | 6 | 89 | 107 |
| ECM22/UF | TATACGA | 0 | 0.3765 | 1 | 0 | 11 | 31 | 165 |
| FKH1 | [GA]TAAA | 7.0449 | 0.0029 | 0.229 | 7 | 4 | 39 | 157 |
| FLO8 | AAAACCT | 0 | 1 | 1 | 0 | 11 | 0 | 196 |
| FZF1 | CGTATCGT | 0 | 1 | 1 | 0 | 11 | 0 | 196 |

**Supplementary Table S4:** Tab separated file containing the genomicfeatures tested foroverrepresentation between circular DNA classified as lost and circular DNA classified as retained (class I and II). Ordered from left to right, the columns contain the following information: 1, genomicfeature tested; 2, Fisher’s exact test odds ratio; 3, Fisher’s exact test *p-value*; 4, adjusted *p-value*; 5, numberof circular DNA containing the genomicfeature in the retained group; 6, numberof circular DNA not containing the genomicfeature i n the retained group; 7, numberof circular DNA containing the genomicfeature in the lost group and 8, number of circular DNA not containing the genomicfeature in the lost group.

| Gene | Motif_reg | OddsRatio | P-value | P-Adjust | In_Retained | Not_in_Retained | In_lost | Not_in_lost |
| --- | --- | --- | --- | --- | --- | --- | --- | --- |
| OriDB | . | 9.2121 | 0.0009 | 0.0062 | 8 | 3 | 44 | 152 |
| centromere | . | NaN | 1 | 1 | 0 | 0 | 0 | 0 |
| long_term | . | 1.4077 | 0.5463 | 0.956 | 1 | 10 | 13 | 183 |
| telomere | . | 43.3333 | 0.0075 | 0.0263 | 2 | 9 | 1 | 195 |
| LTR_retrotransposon | . | 2.35 | 0.3944 | 0.9202 | 1 | 10 | 8 | 188 |
| intron | . | 0.9889 | 1 | 1 | 1 | 10 | 18 | 178 |
| Repetitive | . | 0 | 0.013 | 0.0907 | 3 | 193 | 6 | 5 |

**Supplementary Table S5:**
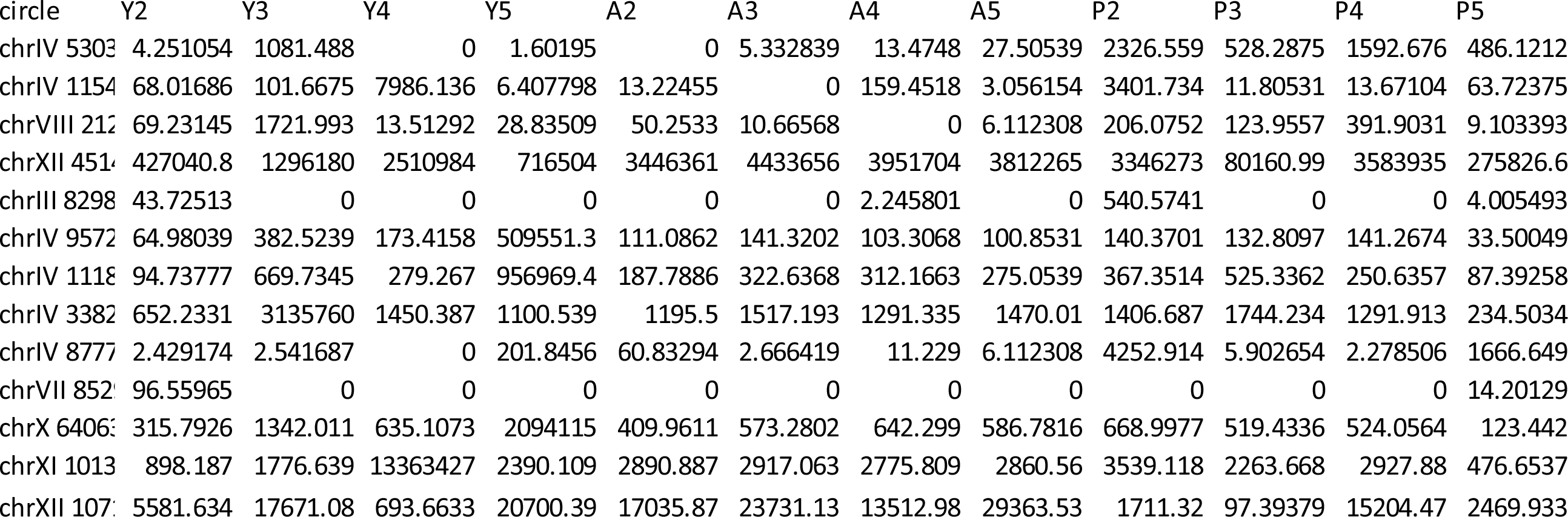
Tab separated file containing the median of the ratios normalized values forthe circular DNA detected in at least two samples of the Mother Enrichment Program experiment. Ordered from left to right, the columns contain the following information: 1, circular DNA detection coordinates togetherwith the numberof samples it was detected; 2-13, sequencing samples (Y indicates the young population, A indicates the aged population and P indicates the progeny population).

**Supplementary Table S6:** Tab separated file containing the plasmid normalized read counts for circular DNA formed from the rDNA locus. Ordered from left to right, the columns contain the following information: 1, sample (Y indicates young and A indicates aged); 2, plasmid normalized read count values; 3: phenotype of the sample.

| sample | rdna | phenotype |
| --- | --- | --- |
| Y10 | 3.4244 | young |
| Y6 | 0.2659 | young |
| Y7 | 23.016 | young |
| Y9 | 1.6526 | young |
| A10 | 68.576 | aged |
| A6 | 491.19 | aged |
| A7 | 142.11 | aged |
| A9 | 1395.7 | aged |

**Supplementary Table S7:** Tab separated file containing the plasmid normalized read counts for circular DNA formed from the Y’ prime telomeric locus. Ordered from left to right, the columns contain the following information: 1, sample (Y indicates young and A indicates aged); 2, plasmid normalized read count values; 3: phenotype of the sample.

| sample | y_prime | phenotype |
| --- | --- | --- |
| Y10 | 0.003225 | young |
| Y6 | 2.89E-05 | young |
| Y7 | 0.023299 | young |
| Y9 | 8.70E-05 | young |
| A10 | 0.046857 | aged |
| A6 | 0.494467 | aged |
| A7 | 0.127192 | aged |
| A9 | 1.116935 | aged |

**Supplementary Table S8:**
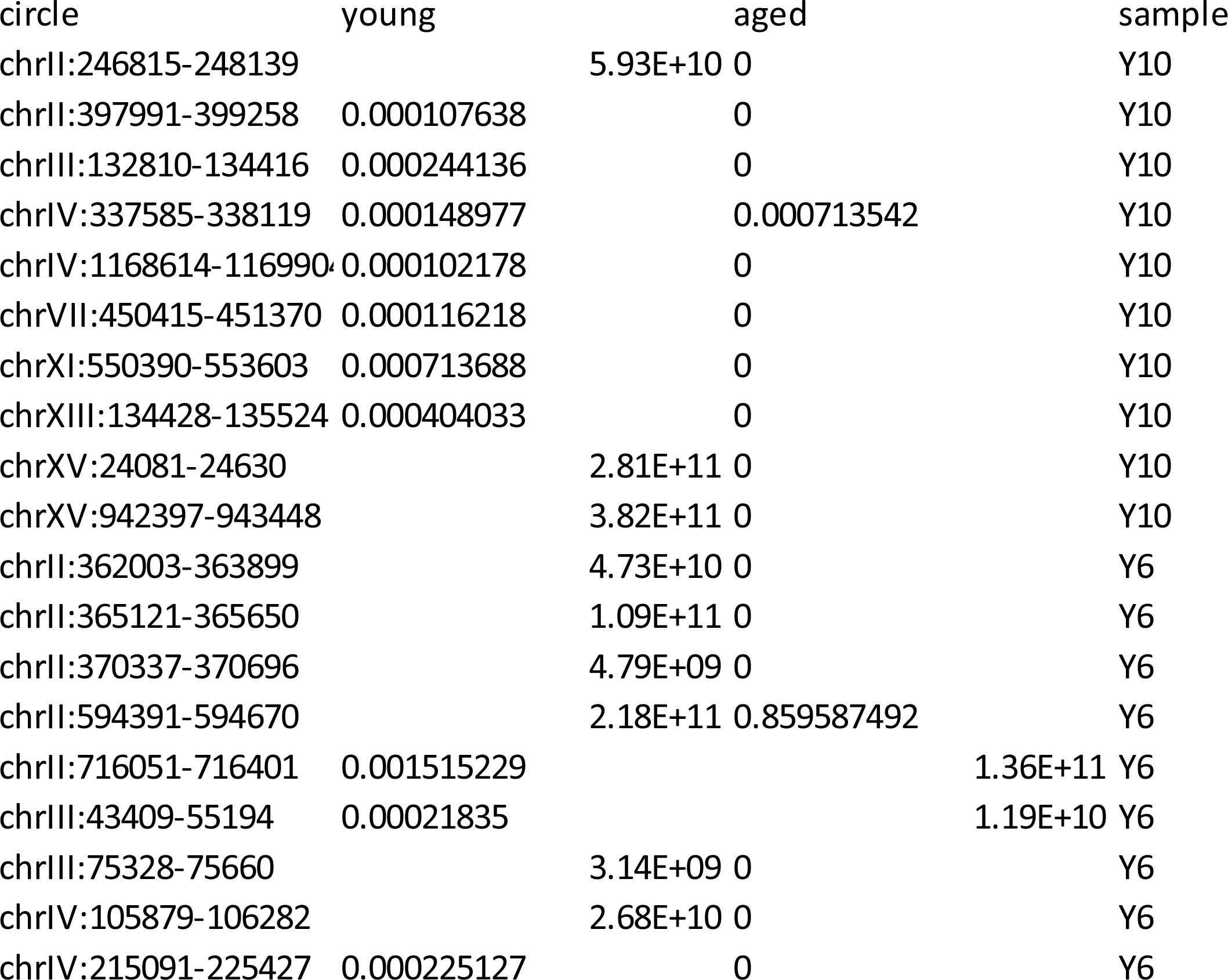
Top 20 lines shown. Tab separated file containing the plasmid normalized read counts for all the circular DNA from other parts of the genome, excluding the rDNA and the Y‘ telomeric locus. Ordered from left to right, the columns contain the following information: 1, circular DNA detection chromosomal coordinates; 2, plasmid normalized read count values in the young sample; 3, plasmid normalized read count values in the aged sample, Young sample where the circular DNA was detected.

## Supplementary files

**Supplementary file S1.** FASTA formatted file containing the DNA sequence of the circular DNA formed from the [*Y’ telomeric ^circle^*] locus.

